# Endocytosis of the damage-associated molecular pattern receptor PEPR1 is BAK1-dependent

**DOI:** 10.1101/2025.03.15.643409

**Authors:** Lucas Alves Neubus Claus, Fausto Andrés Ortiz-Morea, Shao-Li Yang, Shweta Yekondi, Xiangyu Xu, In-Cheol Yeo, Isabelle Vanhoutte, Nemanja Vukašinović, Qian Ma, Ive De Smet, Ping He, Libo Shan, Eugenia Russinova

## Abstract

After cellular damage caused by wounding or pathogens, *Arabidopsis thaliana* endogenous elicitor peptides (Peps) are released into the apoplast, enhancing innate immunity by directly binding to the membrane-localized leucine-rich repeat receptor kinase PEP RECEPTOR1 (PEPR1). Ligand binding induces PEPR1 heterodimerization with the co-receptor BRASSINOSTEROID INSENSITIVE1-ASSOCIATED KINASE1 (BAK1), followed by PEPR1 internalization, both essential for a subset of Pep1-induced responses. However, the role of BAK1 in Pep1-triggered PEPR1 endocytosis remains unclear. Here, we show that the ligand-induced PEPR1 endocytosis depends on its kinase activity and requires BAK1 C-terminal tail phosphorylation, which is equally indispensable for immune signaling and BAK1 internalization. Using a GFP insertional mutagenesis approach, we generated a partially functional GFP-tagged BAK1 to demonstrate that, following Pep1 elicitation, BAK1 and PEPR1 are endocytosed together with similar dynamics. Our findings identify the BAK1 function as a prerequisite for PEPR1 internalization.

## INTRODUCTION

Damage-associated molecular patterns (DAMPs) are endogenous elicitor molecules released into the apoplast after cell damage to trigger immune responses. An example of DAMPs in plants is endogenous plant elicitor peptides (Peps) (Hou et al., 2019). Peps are 23-amino-acid peptides that mature from a conserved C-terminal region of larger precursor proteins, designated PROPEPs (Yamaguchi and Huffaker, 2011), which belong to a family of eight members (PROPEP1 to PROPEP8) in *Arabidopsis thaliana* (Arabidopsis). Peps are recognized by two homologous pattern recognition receptors (PRRs), the leucine-rich repeat (LRR) receptor kinases (RKs) PEP RECEPTOR1 (PEPR1) and PEPR2 (Krol et al., 2010; Yamaguchi et al., 2010; Bartels et al., 2013). The Pep-PEPR ligand-receptor pair is also present in other dicots, such as *Solanaceae* and *Fabaceae* (Trivilin et al., 2014; Bartels and Boller, 2015), as well as monocots, such as *Zea mays* (maize) (Huffaker et al., 2011, 2013).

Upon Pep1 binding, PEPR1 associates with the LRR co-receptor, BRASSINOSTEROID-INSENSITIVE1 (BRI1)-ASSOCIATED RECEPTOR KINASE1 (BAK1)/SOMATIC EMBRYOGENESIS RECEPTOR KINASE (SERK3), hereafter referred to as BAK1 (Tang et al., 2015; Yamada et al., 2016). BAK1 is a promiscuous protein that regulates multiple signaling pathways controlling growth, cell death, and innate immunity by forming ligand-dependent complexes with several plasma membrane (PM)-localized RKs (Schulze et al., 2010; Schwessinger et al., 2011). This raises the question of how BAK1 distinguishes between different signaling pathways and responds accordingly. BAK1 has been shown to discriminate between immune and brassinosteroid (BR) hormone responses in a phosphorylation-dependent manner (Perraki et al., 2018). Accordingly, the *bak1-5* mutant allele, which carries a single amino acid substitution (Cys to Tyr at position 408) in the kinase domain, leading to reduced phosphorylation, is severely impaired in immune responses, but not in responses to BRs or cell death (Schwessinger et al., 2011).

After perceiving Pep1, PEPR1 undergoes clathrin-mediated endocytosis (CME) (Mbengue et al., 2016; Ortiz-Morea et al., 2016), an evolutionarily conserved process that is essential for immunity and development in plants (Claus et al., 2018). CME regulates the abundance of signaling components in the PM that are important for nutrient uptake (Takano et al., 2010; Wang et al., 2017; Dubeaux et al., 2018), hormone signaling (Irani et al., 2012), pathogen recognition, and stress responses (Krol et al., 2010; Choi et al., 2014). Previously, PEPR1 was found to be internalized from the PM to the late endosome/multivesicular bodies (LE/MVBs), bypassing the trans-Golgi network/early endosomes (TGN/EEs), from where it is targeted to the vacuole for degradation (Ortiz-Morea et al, 2016). PEPR1 shares an endomembrane trafficking pathway with the PRRs recognizing the microbe-associated molecular patterns (MAMPs), such as FLAGELLIN-SENSING2 (FLS2) and EF-TU RECEPTOR (EFR) (Robatzek et al., 2006; Mbengue et al., 2016), but differs from the well-studied BR receptor BRI1, which is internalized from the PM to TGN/EE, where it can be recycled back to the PM or targeted to the vacuole for degradation (Irani et al., 2012).

Endocytosis is often considered as a mechanism for removing receptors from the PM to avoid continuous signaling that is potentially harmful to cells (Sigismund et al., 2012; Jaillais and Vert, 2016). For example, in plants, blocking BRI1 endocytosis at the PM results in constitutive BR responses, indicating that endocytosis attenuates BR signaling (Irani et al., 2012; Martins et al., 2015; Zhou et al., 2018; Liu et al., 2020). However, disruption of PRR endocytosis affects negatively plant defense responses (Ortiz-Morea et al., 2016; Mbengue et al., 2016; Ekanayake et al., 2021).

Although considerable progress has been made in understanding the mechanisms of plant receptor endocytosis, they remain poorly understood compared to their mammalian counterparts. Post-translational modifications (PTMs), such as phosphorylation and ubiquitination, are important regulators of FLS2 and BRI1 receptor endocytosis (Robatzek et al., 2006; Mbengue et al., 2016; Lu et al., 2011; Martins et al., 2015; Zhou et al., 2018; Ma et al., 2020). How the co-receptor BAK1 regulates endocytosis of the major receptors is still unclear, because studies of BAK1 endocytosis in Arabidopsis have been hampered by the observation that the C-terminally tagged BAK1 is not fully functional (Ntoukakis et al., 2011; Lozano-Durán et al., 2013).

Here we sought to investigate the role of BAK1 in PEPR1 endocytosis. We found that, in contrast to BRI1, the Pep1-induced endocytosis of PEPR1 requires BAK1 phosphorylation in the C-terminal tail domain that is essential for both DAMP signaling and BAK1 endocytosis. We further showed that the kinase activity of PEPR1 is necessary for its internalization and that PEPR1 but not BAK1 was able to trans-phosphorylate *in vitro*. A GFP insertional mutagenesis approach was used to generate a partially functional GFP-tagged BAK1, showing that following Pep1 application BAK1 and PEPR1 are endocytosed together.

## RESULTS

### Pep1-triggered responses are impaired by the C-terminal GFP-tagged BAK1 fusion protein

Arabidopsis plants expressing C-terminally tagged BAK1 variants exhibit impaired immune responses to flg22 and elf18, but largely retain normal responses to exogenous BRs (Ntoukakis et al., 2011; Lozano-Durán et al., 2013). Therefore, we tested whether the C-terminal tag of BAK1 also interferes with responses to exogenous Pep1. To this end, we generated transgenic Arabidopsis plants expressing native promoter-driven BAK1-GFP in the *bak1-4* mutant and analyzed two independent transgenic lines (Supplemental Figure 1A-1E). BAK1-GFP complemented the growth phenotype of *bak1-4* and remained functional in BR responses (Supplemental Figure 1A-1E), although it caused slight BR hypersensitivity, possibly due to increased expression levels (Supplemental Figure 1B and 1C). As previously described, exogenous Pep1 inhibited wild-type root growth in a dose-dependent manner, whereas *bak1-4* was hypersensitive to Pep1 (Figure 1A and 1B) due to sensitized PEPR signaling towards cell death (Yamada et al., 2016). However, BAK1-GFP/*bak1-4* plants were insensitive to the Pep1-induced root growth arrest, both MAPK activation and reactive oxygen species (ROS) burst triggered by Pep1 were lower than in the wild-type, although the MAPK activation was less affected (Figure 1C and 1D). Our findings indicate that, in addition to flg22 and elf18, the C-terminal tag of BAK1 impairs Pep1 responses.

**Figure 1.**
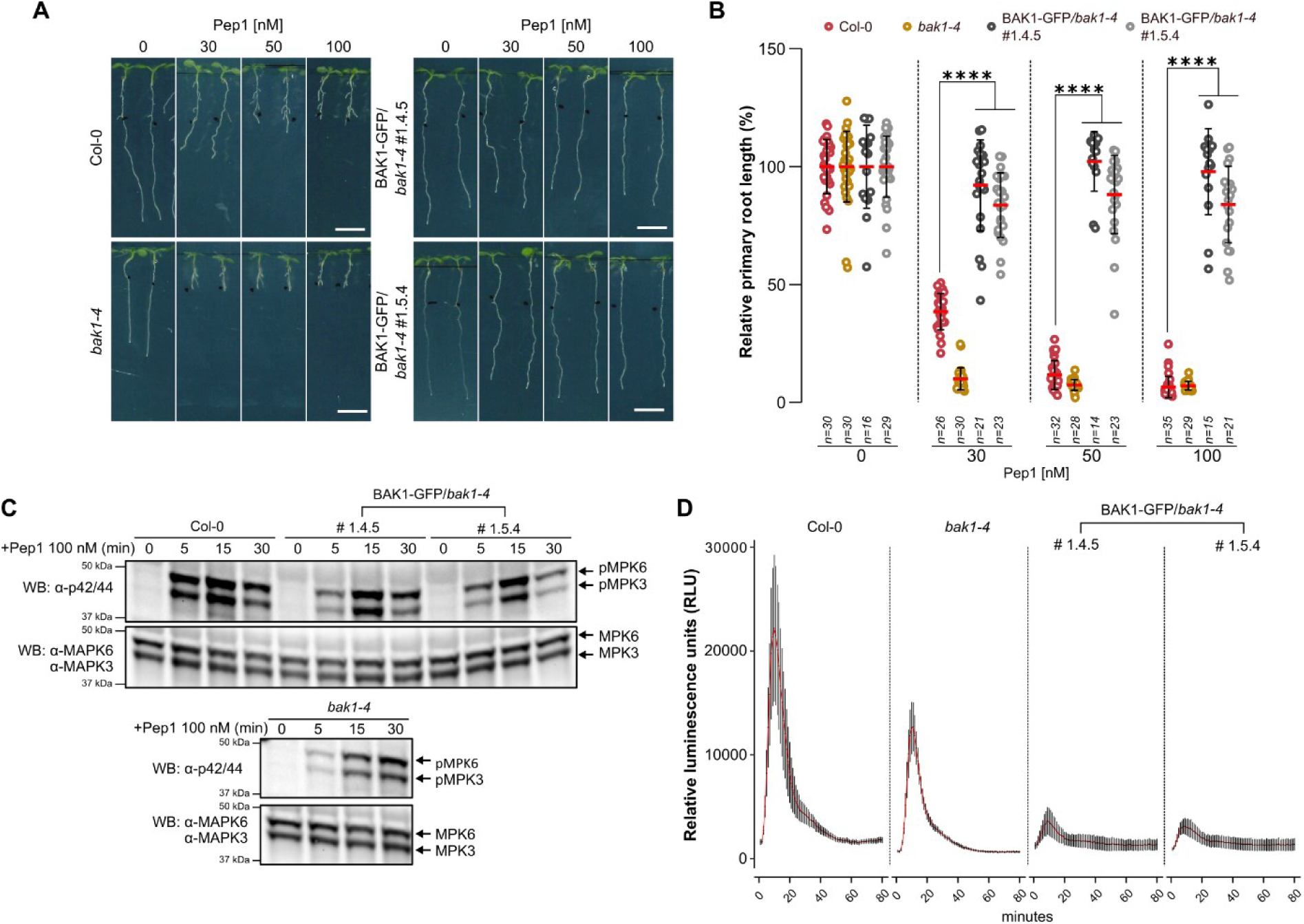
C-terminally tagged BAK1-GFP is not fully functional in Pep1-induced immune responses. **(A)** Representative images of root growth inhibition caused by different Pep1 concentrations. Seedlings of wild-type Col-0, *bak1-4*, and two independent *pBAK1-BAK1-GFP*/*bak1-4* transgenic lines (#1.4.5 and #1.5.4) were grown for 4 days on ½MS agar medium and transferred to ½MS agar medium supplemented with different concentrations of Pep1 or water (mock) for a further 3 days. Scale bars, 10 mm. **(B)** Primary root length of seedlings in **(A)** presented as relative to the mock-treated control for each genotype. Individual measurements, average, and SD are represented by dots, red lines, and whiskers, respectively. Two-way ANOVA followed by Tukey’s test was used to determine significant differences.\*\*\*\**P<*0.0001*; n*, number of seedlings analyzed. **(C)** MAPK activation assay of seven-day-old seedlings in **(A)** treated with 100 nM Pep1 for the indicated time points. Phosphorylated MPK3 (pMPK3) and MPK6 (pMPK6) were detected by Western Blotting (WB) with α-p42/44 antibody (top). Protein loading is shown by total MPK3 and MPK6 detected with α-MPK3 and α-MPK6 antibodies (bottom). **(D)** Pep1-induced reactive oxygen species (ROS) production. Eight leaf discs from four 5-week-old plants in (A) treated with 100 nM of Pep1. ROS production measured as RLU was detected at the indicated time points. Red lines indicate means ± SD.

### BAK1 differentially regulates endocytosis of PEPR1 and BRI1

Since exogenous Pep1 induces PEPR1 endocytosis (Ortiz-Morea et al., 2016), we tested whether the C-terminally GFP-tagged BAK1 affects endocytosis of the Pep1-bound PEPR1 by monitoring the internalization of the bioactive TAMRA-labelled Pep1 (TAMRA-Pep1) (Ortiz-Morea et al., 2016) in BAK1-GFP/*bak1-4* plants (Figure 2A and 2B). As a control, we quantified endocytosis of the fluorescent Alexa Fluor 647-castasterone (AFCS), a bioactive fluorescently labelled BR analogue that is internalized together with BRI1 (Irani et al., 2012) (Figure 2C and 2D). In contrast to the wild-type, TAMRA-Pep1 internalization was strongly inhibited in BAK1-GFP/*bak1-4* plants, with few TAMRA-Pep1 puncta appearing inside the cell and a strong signal remaining in the PM (Figure 2A and B). However, TAMRA-Pep1 internalization in the *bak1-4* mutant was not significantly different from that of the wild-type at the analyzed time point (40 min chase) (Figure 2A and 2B). Consistently, exogenous Pep1-induced PEPR1-GFP internalization in *bak1-4* and *pepr1pepr2* at 40 min (Supplemental Figure 2A) resembled TAMRA-Pep1 (Figure 2A and 2B) in both mutants but was partially inhibited at later time points (60-120 min) in *bak1-4*. The AFCS uptake in BAK1-GFP/*bak1-4* and *bak1-4* plants was comparable to or slightly higher than that of the wild-type (Figure 2C and 2D), suggesting that C-terminally tagged BAK1 affects Pep1-PEPR1, but not AFCS-BRI1 endocytosis.

**Figure 2.**
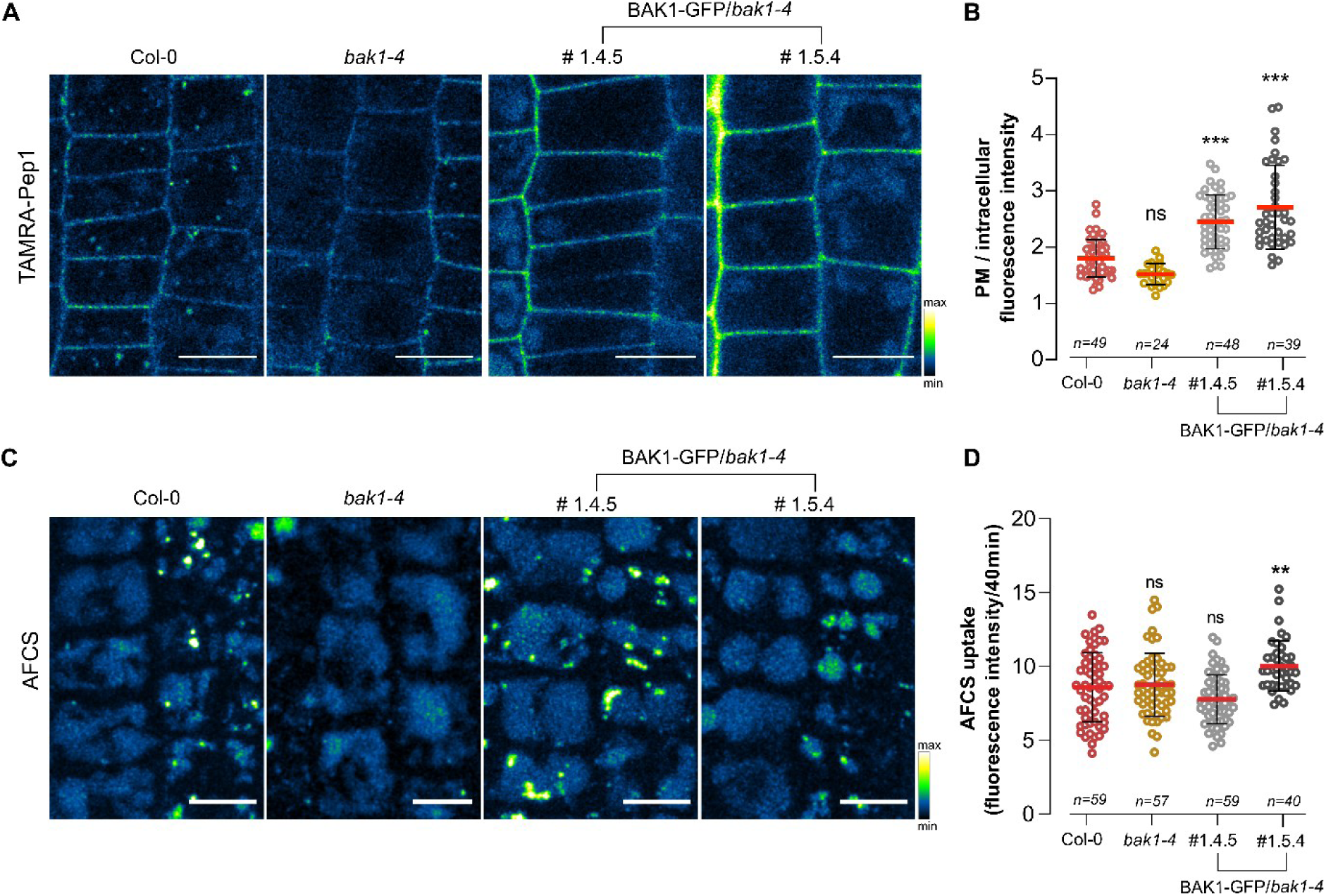
Pep1-triggered PEPR1 endocytosis requires functional BAK1. **(A, C)** Representative images of TAMRA-Pep1 **(A)** and AFCS **(C)** root uptake visualized by color-based fluorescence intensity coding. Five-day-old seedlings of wild-type Col-0, *bak1-4*, and two independent *pBAK1-BAK1-GFP*/*bak1-4* transgenic lines were treated with 100 nM TAMRA-Pep1 for 10 s (pulse) and with 40 µM AFCS for 20 min (pulse), respectively, and imaged after 40 min (chase). Epidermal cells of root meristematic zone were imaged. Scale bars, 10 µm. **(B, D)** Fluorescence intensity measurements of plasma membrane (PM) vs intracellular compartments of the **(B)** TAMRA-Pep1 uptake in **(A)** and **(D)** the AFCS uptake after 40 min chase in **(C)**. Individual measurements, average, and SD are represented by dots, red line, and whiskers, respectively. One-way ANOVA followed by Tukey’s test was used to determine significant differences.\*\*\**P*<0.001, \*\**P*<0.01; *n*, number of cell analyzed; ns, not significant.

To confirm the importance of BAK1 for PEPR1 endocytosis, we examined the internalization of PEPR1-GFP and TAMRA-Pep1 in different *serk* mutant combinations after 40 min chase (Supplemental Figure 2B-2E). TAMRA-Pep1 internalization was unaffected in the *bak1-3serk4-1* double and the *serk1-1serk2-1serk4-1* triple mutant but was strongly inhibited in the *serk1-1bak1-4* double mutant and the *serk1-1serk2-1bak1-4* triple mutant (Supplemental Figure 2B-2E). These findings suggest that BAK1 and SERK1 redundantly regulate PEPR1 endocytosis.

Taken together, our results show that C-terminally tagged BAK1 impairs ligand-induced endocytosis of PEPR1 but not of BRI1, suggesting that BAK1 may regulate the internalization of BR-bound BRI1 and Pep1-bound PEPR1 differently.

### The C-terminal phosphorylation of BAK1 is essential for PEPR1 endocytosis and signaling

Endocytosis of membrane proteins requires prior phosphorylation (Wang et al., 2017; Zhou et al., 2018). The phosphorylation relationship between PEPR1 and BAK1 has not been previously investigated. In vitro, the cytoplasmic domains (CDs) of both proteins displayed autophosphorylation activity, although PEPR1 activity was weaker, whereas their respective kinase-dead mutants (Km) were inactive (Supplemental Figure 3A and Supplemental Dataset 1). Notably, PEPR1-CD trans-phosphorylated BAK1-Km-CD, whereas BAK1-CD did not trans-phosphorylate PEPR1-Km-CD, despite retaining the ability to trans-phosphorylate BOTRYTIS-INDUCED KINASE 1 (BIK1)-Km (Lu et al., 2010) (Supplemental Figure 3B). Mass spectrometry (MS) identified six PEPR1-mediated phosphorylation sites on BAK1 (S290, T324, T446, T449, T450, and T455), four of which (T446, T449, T450, and T455) are located in the activation loop (Supplemental Dataset 1). Furthermore, PEPR1-CD, but not PEPR1-Km-CD, reduced BAK1 autophosphorylation *in vitro*, while the PEPR1-CD activity appeared enhanced by BAK1-CD (Supplemental Figure 3A). A C-terminal cluster of BAK1 autophosphorylation sites (S602, T603, S604, and S612) is essential for a subset of its functions (Perraki et al., 2018; Bender et al., 2021). Because, C-terminal GFP-tagged BAK1 loses phosphorylation at one of these sites (S612) (Perraki et al., 2018), we hypothesized that this cluster is critical for Pep1-PEPR1 endocytosis and signaling. To test this, we compared TAMRA-Pep1 uptake in the phosphorylation-deficient BAK1^S612A^ and BAK1^S602AT603AS604A^ (hereafter BAK1^AAA^) mutants with wild-type. Both mutants showed markedly reduced uptake, with a more pronounced effect in BAK1^S612A^ (Figure 3A and 3B). Similarly, of PEPR1-GFP endocytosis was impaired in both phosphorylation-deficient BAK1 variants expressed in the *bak1-4* background, again with a stronger reduction in the BAK1^S612A^ mutant (Figure 3C and 3D).

**Figure 3.**
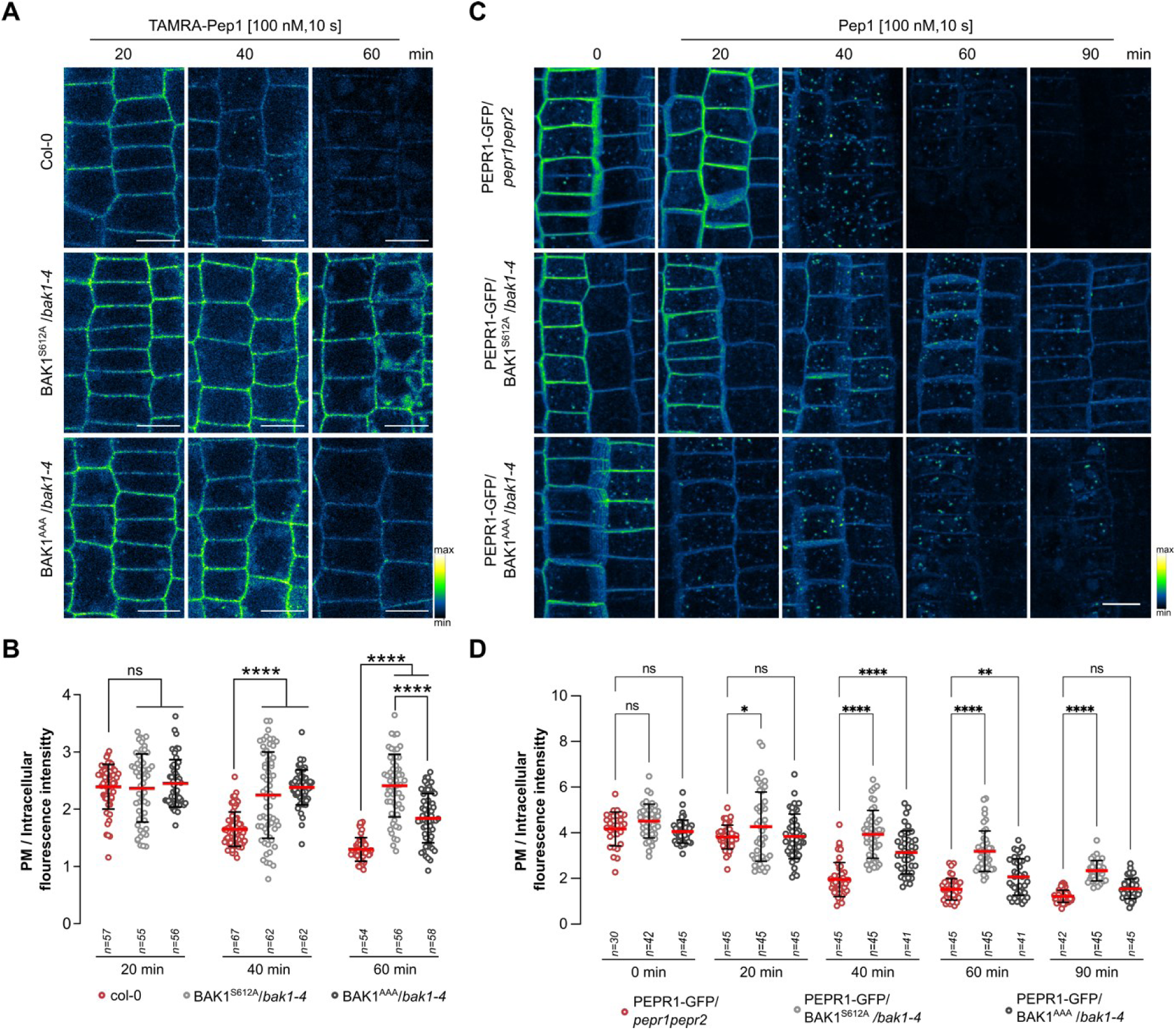
The BAK1 phosphorylation essential for flagellin-induced signaling influences PEPR1 endocytosis. **(A)** Representative images of TAMRA-Pep1 root uptake visualized by color-based fluorescence intensity coding. Five-day-old wild-type Col-0, *pBAK1-BAK1^S612A^*/*bak1-4* and *pBAK1-BAK1^AAA^*/*bak1-4* seedlings were treated with 100 nM TAMRA-Pep1 for 10 s (pulse) and imaged at indicated time points (chase). Epidermal cells of root meristematic zone were imaged. Scale bars, 10 µm. **(B)** Plasma membrane (PM) vs. intracellular fluorescence intensity measurements of the TAMRA-Pep1 uptake in **(A)**. Individual measurements, average, and SD are indicated as dots, red line, and whiskers, respectively. One-way ANOVA followed by Tukey’s test was used to determine significant differences.\*\*\*\**P<*0.0001. **(C)** Representative images of PEPR1-GFP endocytosis visualized by color-based fluorescence intensity coding. Five-day-old *pRPS5A-PEPR1-GFP*/*pepr1pepr2*, *pRPS5A-PEPR1-GFP*/*pBAK1-BAK1^S612A^*/*bak1-4* and *pRPS5A-PEPR1-GFP*/*pBAK1-BAK1^AAA^*/*bak1-4* seedlings were treated with 100 nM Pep1 for 10 s (pulse) and imaged at indicated time points (chase). Images are a maximum intensity Z-projection. Epidermal cells of root meristematic zone were imaged. Scale bar, 10 µm. **(D)** PM vs. intracellular fluorescence intensity measurements of PEPR1-GFP internalization in **(C)**. Dots, red lines, and whiskers represent individual measurements, red lines, and SD, respectively. One-way ANOVA followed by Tukey’s test was used to determine significant differences.\*\*\*\**P*<0.0001, \*\*\**P*<0.001, \*\**P*<0.01, \**P*<0.05; *n*, number of cell analyzed **(B, D)**; ns, not significant.

Since PEPR1-Pep1 endocytosis was impaired in the BAK1^S612A^ and BAK1^AAA^ mutants, we suspected that the Pep1-induced signaling would also be disturbed. To test this hypothesis, we performed root growth, MAPK activation and ROS burst assays after Pep1 treatment (Figure 4; Supplemental Figure 4). Similar to BAK1-GFP/*bak1-4*, BAK1^S612A^/*bak1-4* and BAK1^AAA^/*bak1-4* plants were strongly insensitive to Pep1 treatment (Figure 4A and 4B) and both MAPK activation (Figure 4C) and ROS burst were reduced compared to the wild-type (Figure 4D). To further confirm that BAK1 phosphorylation is crucial for Pep1-induced endocytosis, we analyzed the *bak1-5* mutant, which carries the C408Y point mutation, responsible for the BAK1 hypoactivity (Schwessinger et al., 2011). *bak1-5* resembled BAK1-GFP/*bak1-4*, BAK1^S612A^/*bak1-4*, and BAK1^AAA^/*bak1-4* plants in that it was less responsive to exogenous Pep1 in the root growth inhibition assay (Supplemental Figure 5A and 5B), consistent with previous findings for the MAMPs flg22 and elf18 (Schwessinger et al., *2*011). Additionally, Pep1-PEPR1 endocytosis was impaired in *bak1-5* mutant (Supplemental Figure 5C and 5D). Taken together, our results indicate that C-terminal BAK1 phosphorylation, which is indispensable for flg22, elf18, and Pep1 signaling, is also a prerequisite for Pep1-induced PEPR1 endocytosis.

**Figure 4.**
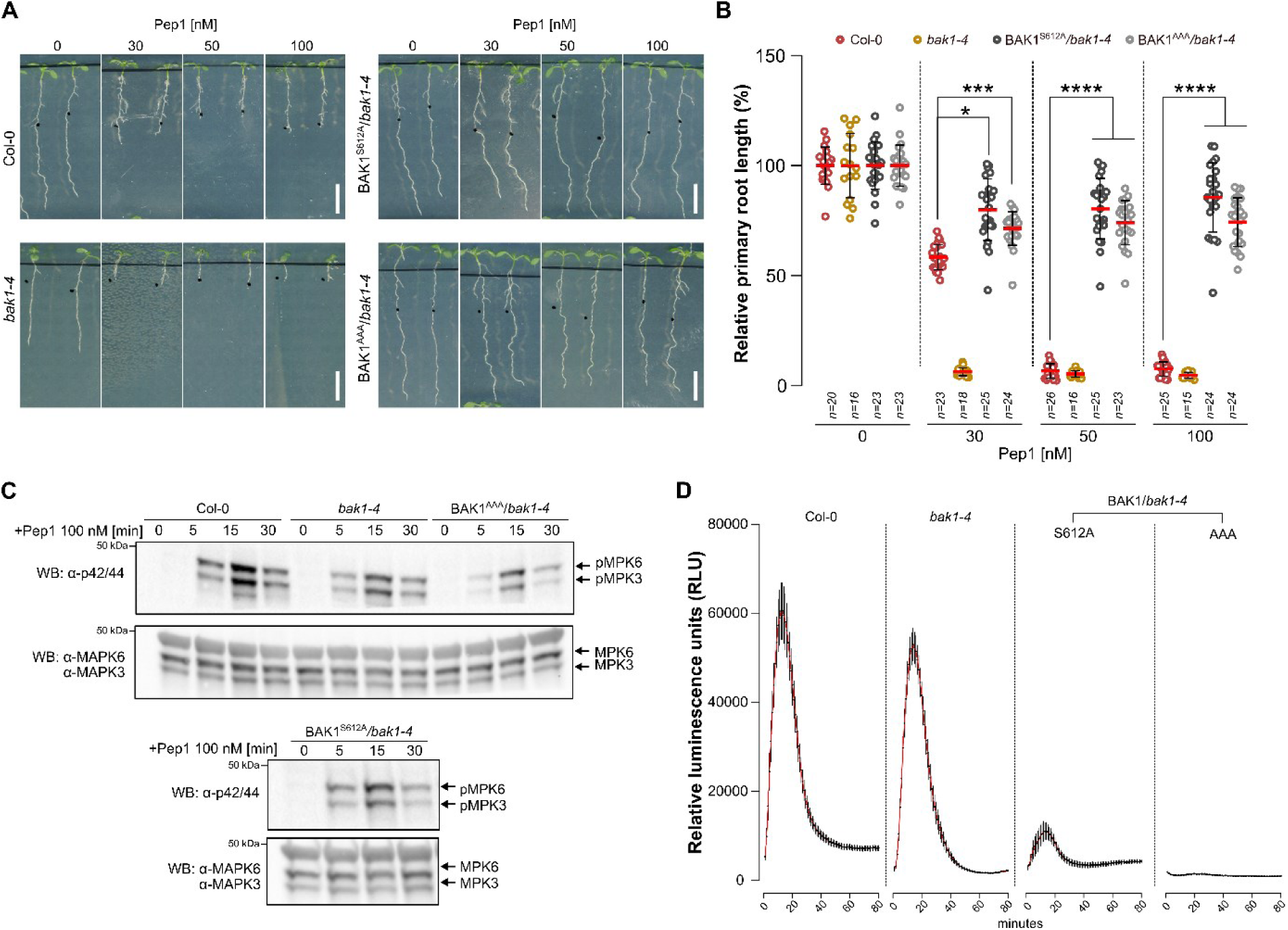
Flagellin-and Pep1-induced signaling pathways require the same BAK1 phosphorylation signature. **(A)** Representative images of root growth inhibition caused by different Pep1 concentrations. Seedlings of wild-type Col-0, *bak1-4*, *pBAK1-BAK1^S612A^*/*bak1-4* and *pBAK1-BAK1^AAA^*/*bak1-4* were grown for 4 days on ½MS agar plates and transferred to ½MS solid media supplemented with different concentrations of Pep1 or water (mock) for a further 3 days. Scale bars, 10 mm. **(B)** Primary root length of seedlings in **(A)** presented as relative to the mock-treated control for each genotype. Dots, red lines, and whiskers represent individual measurements, averages, and SD, respectively. The asterisk indicates statistically significant difference analyzed with two-way ANOVA followed by Tukey’s test. \*\*\*\**P*<0.0001, \*\*\**P*<0.001, \**P*<0.05; *n*, number of seedlings analyzed; ns, not significant. **(C)** MAPK activation assay of 5-day-old seedlings in **(A)** treated with 100 nM Pep1 for the indicated time points. Phosphorylated MPK3 (pMPK3) and MPK6 (pMPK6) were detected by Western Blotting (WB) with the α-p42/44 antibody (top). Protein loading is shown by total MPK3 and MPK6 detected with the α-MPK3 and α-MPK6 antibodies (bottom). **(D)** Pep1-induced reactive oxygen species (ROS) production. Eight leaf discs from four 5-week-old plants in **(A)** were treated with 100 nM of Pep1. ROS production measured as RLU was detected at the indicated time points. Red lines indicate means ± SD.

### Pep1 endocytosis requires kinase-active PEPR1

To investigate whether phosphorylation of PEPR1 after ligand binding is required for its internalization, we generated transgenic plants expressing a kinase-dead PEPR1-GFP (PEPR1-Km-GFP) in the *pepr1pepr2* double mutant (Figure 5A). As previously reported (Ortiz-Morea et al., 2016), without elicitation, PEPR1-GFP localized mainly to the PM, with a few puncta inside the cell. Pep1 treatment triggered its internalization, with an endocytosis peak after 40 min. However, endocytosis of PEPR1-Km-GFP after Pep1 application was decreased, and the delay was most apparent 90-120 min after elicitation, when the fluorescence signal almost disappeared, although the PM signal remained stronger. Interestingly, without exogenous Pep1, PEPR1-Km-GFP was observed in intracellular puncta, possibly indicating either endocytosis of PEPR1-Km-GFP without ligand binding or increase of its *de novo* production and exocytosis (Figure 5A). To further investigate, we treated PEPR1-GFP and PEPR1-Km-GFP plants with Brefeldin A (BFA), an indirect inhibitor of endosomal trafficking, that induces aggregation of the *trans*-Golgi network/early endosome (TGN/EE) compartments, forming BFA bodies (Geldner et al., 2003) (Figure 5B). Without Pep1 elicitation, PEPR1-GFP was only weakly present in the BFA bodies, but after Pep1 application, PEPR1-GFP puncta were excluded from the BFA bodies and localized to the periphery of the TGN/EE aggregates. Notably, under both Pep1-treated or mock conditions, the PEPR1-Km-GFP labelling of the BFA bodies was strong, suggesting that the BFA bodies accumulate both internalized and secreted PEPR1-Km-GFP (Figure 5B).

**Figure 5.**
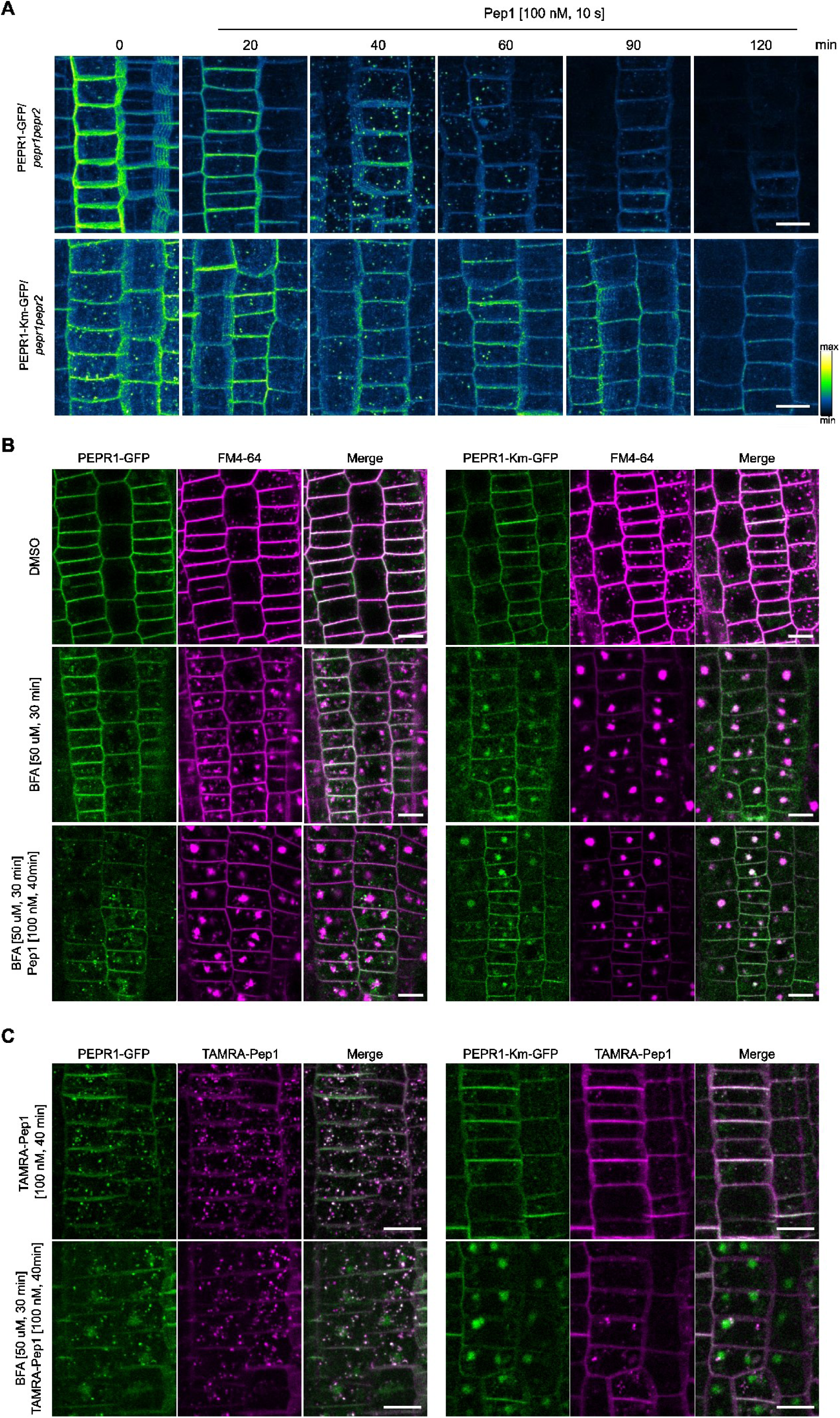
The kinase activity of PEPR1 is required for its endocytosis. **(A)** Representative images of PEPR-GFP and PEPR1-Km-GFP (PEPR1-kinase mutant-GFP) internalization visualized by color-based fluorescence intensity coding. Five-day-old *pRPS5A-PEPR1-GFP*/*pepr1pepr2* and *pRPS5A-PEPR1-Km-GFP*/*pepr1pepr2* seedlings were treated with 100 nM of Pep1 for 10 s (pulse) and imaged at the indicated time points (chase). Epidermal cells of root meristematic zone were imaged. Images are maximum intensity Z-projections. Scale bars, 10 µm. **(B, C)** Representative images of PEPR1-GFP/*pepr1pepr2* and PEPR1-Km-GFP/*pepr1pepr2* internalization in the presence of BFA **(B)** and endocytosis of TAMRA-Pep1 in the absence and presence of BFA **(C)**. Seedlings as in **(A)** were treated with DMSO (mock), BFA (50 µM, 30 min) and Pep1 (100 nM, 40 min) after pre-treatment with BFA (50 µM, 30 min) **(B)**. Seedlings as in **(A)** were treated with TAMRA-Pep1 (100 nM, 40 min) or TAMRA-Pep1 (100 nM, 40 min) after pre-treatment with BFA (50 µM, 30 min) **(C)**. Epidermal cells of the root meristematic zone were imaged. Images are maximum Z-projections. 2 μM FM64 (*N*-(3-triethylammoniumpropyl)-4-(6-(4-(diethylamino) phenyl) hexatrienyl) pyridinium dibromide) dye was used to visualize general endocytosis in **(B)**. Scale bars, 10 µm.

Cycloheximide (CHX), which blocks *de novo* protein synthesis, is routinely used to avoid the interference from secreted proteins when analyzing endocytosis or recycling (Liu et al., 2020). Unexpectedly, CHX blocked PEPR1 internalization (Supplemental Figure 6A) and disturbed the Pep1-triggered MAPK activation (Supplemental Figure 6B). To overcome this limitation, we used TAMRA-Pep1, which allows the tracking of PEPR1 internalization from the PM. In contrast to PEPR1-GFP, the endocytosis of PEPR1-Km-GFP was reduced after TAMRA-Pep1 application (Figure 5C). After BFA treatment, TAMRA-Pep1 does not label the BFA bodies in plants expressing either PEPR1-GFP or PEPR1-Km-GFP, indicating that the accumulated PEPR1-Km-GFP signal in the BFA bodies is mainly due to newly protein synthesis and exocytosis (Figure 5C). Thus, PEPR1 kinase activity is required for its endocytosis.

### BAK1 and PEPR1 are co-internalized

To investigate whether BAK1 is internalized together with PEPR1 we used an insertional GFP mutagenesis approach to generate a functional BAK1-GFP fusion by inserting the GFP at different positions into the BAK1 protein (Supplemental Figure 7A and 7B). All constructs were introduced into the *bak1-4* null mutant under the control of the *BAK1* promoter and several independent transgenic lines of each construct were analyzed. Two constructs with GFP insertions into BAK1 between the juxta membrane and the kinase domains (BAK1K253-GFP and BAK1Q273-GFP) and two constructs in which the GFP was inserted into the C-terminal tail of BAK1 (BAK1M580-GFP and BAK1A593-GFP) were selected (Supplemental Figures 1A-1C and 8A-8B). Of all transgenic lines initially tested in a Pep1-induced root growth inhibition assay, only BAK1K253-GFP and BAK1Q273-GFP partially recovered from the Pep1-triggered root inhibition, although they remained more sensitive to Pep1 than plants expressing the C-terminally tagged BAK1. This phenotype was confirmed with different Pep1 concentrations (Figure 6A and 6B and Supplemental Figure 8C-8E). However, Pep1-triggered MAPK activation in BAK1K253-GFP and BAK1Q273-GFP plants was similar to that of plants expressing BAK1-GFP (Figure 6C). The two mutant lines were more responsive to Pep1 application in terms of ROS burst, although they did not produce the ROS levels of the wild-type (Figure 6D). We further examined the responses of these lines to exogenous BRs by assessing the hypocotyl elongation in the dark and the accumulation of dephosphorylated BES1 (dBES1), which serves as a readout for BR signaling activation (Yin et al., 2002) (Supplemental Figure 1F and 1G). The hypocotyl growth of BAK1K253-GFP and BAK1Q273-GFP seedlings was sensitive to 100 nM, but not 10 nM brassinolide (BL) and the BES1 dephosphorylation levels were similar to those of the wild-type and BAK-GFP expressing plants, suggesting that the response to exogenous BRs in these lines was not notably affected by the GFP tag.

**Figure 6.**
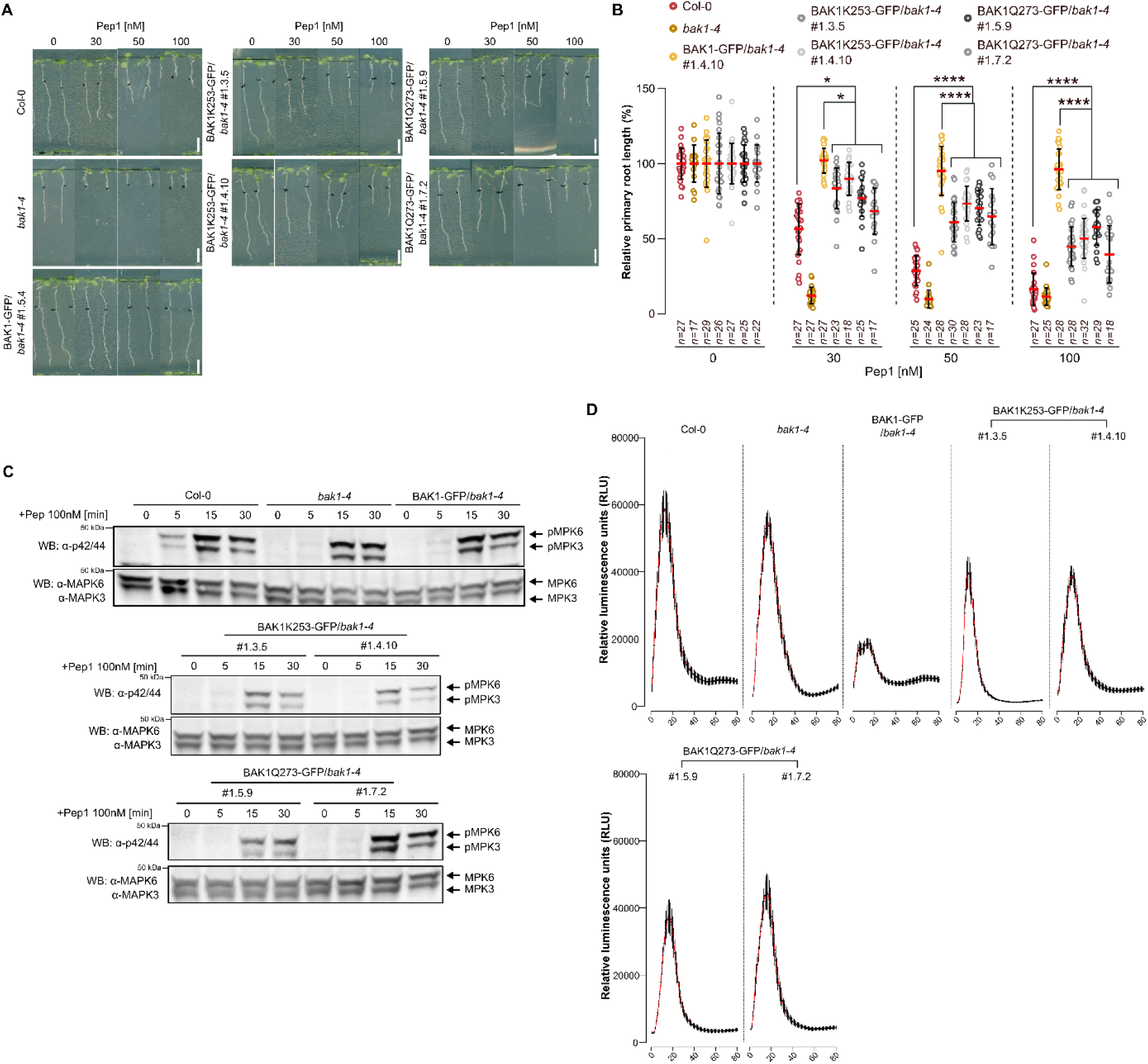
Arabidopsis plants expressing BAK1K253-GFP and BAK1Q273-GFP are partially sensitive to exogenous Pep1. **(A)** Representative images of root growth inhibition caused by different Pep1 concentrations. Seedlings of wild-type Col-0, *bak1-4*, *pBAK1-BAK1-GFP*/*bak1-4*, and two independent transgenic lines of each *pBAK1-BAK1K253-GFP*/*bak1-4* and *pBAK1-BAK1Q273-GFP/bak1-4* were grown for 4 days on ½MS agar plates and then transferred to a ½MS solid media supplemented with different concentrations of Pep1 or water (mock) for a further 3 days. Scale bars, 10 mm. **(B)** Primary root length of seedlings in **(A)** presented as relative to the mock-treated control for each genotype. Dots, red lines, and whiskers represent individual measurements, averages, and SD. Statistically significant difference analyzed with two-way ANOVA followed by Tukey’s test; \*\*\*\**P*<0.0001, \**P*<0.05; *n*, number of seedlings analyzed. **(C)** MAPK activation assay after Pep1 application of 5-day-old seedlings as in **(A)** treated with 100 nM Pep1 for the indicated time points. Phosphorylated MPK3 (pMPK3) and MPK6 (pMPK6) were detected by Western Blotting (WB) with the α-p42/44 antibody (top). Protein loading is shown by total MPK3 and MPK6 detected with the α-MPK3 and α-MPK6 antibodies (bottom). **(D)** Pep1-induced reactive oxygen species (ROS) production. Eight leaf discs from four 5-week-old plants in (A) were treated with 100 nM of Pep1. ROS production measured as RLU was detected at the indicated time points. Red lines indicate mean ± SD.

Next, we took advantage of the partially functional BAK1K253-GFP and BAK1Q273-GFP constructs to analyze TAMRA-Pep1 uptake in the corresponding Arabidopsis transgenic plants (Figure 7A and 7B). Interestingly, the endocytosis of TAMRA-Pep1 was restored in BAK1K253-GFP and BAK1Q273-GFP plants compared to BAK1-GFP (Figure 7A and 7B). After a 40-minute chase, TAMRA-Pep1 accumulated in the PM of plants expressing BAK1-GFP, whereas it was clearly internalized in plants expressing BAK1K253-GFP and BAK1Q273-GFP. Based on the recovery of PEPR1 endocytosis in the BAK1K253-GFP and BAK1Q273-GFP lines, we hypothesized that PEPR1 and BAK1 are endocytosed together. Therefore, we analyzed the co-localization between BAK1-GFP and TAMRA-Pep1 (Figure 7C and 7D). Both BAK1K253-GFP and BAK1Q273-GFP co-localized with the TAMRA-Pep1 after a 40-minute chase, though not all TAMRA-Pep1 puncta contained the BAK1-GFP (Figure 7C). Furthermore, AFCS uptake experiments confirmed that BRI1 endocytosis was not impaired in the BAK1K253-GFP and BAK1Q273-GFP lines (Figure 7E and 7F), consistent with our observations that BR signaling remained unaffected in these plants.

**Figure 7.**
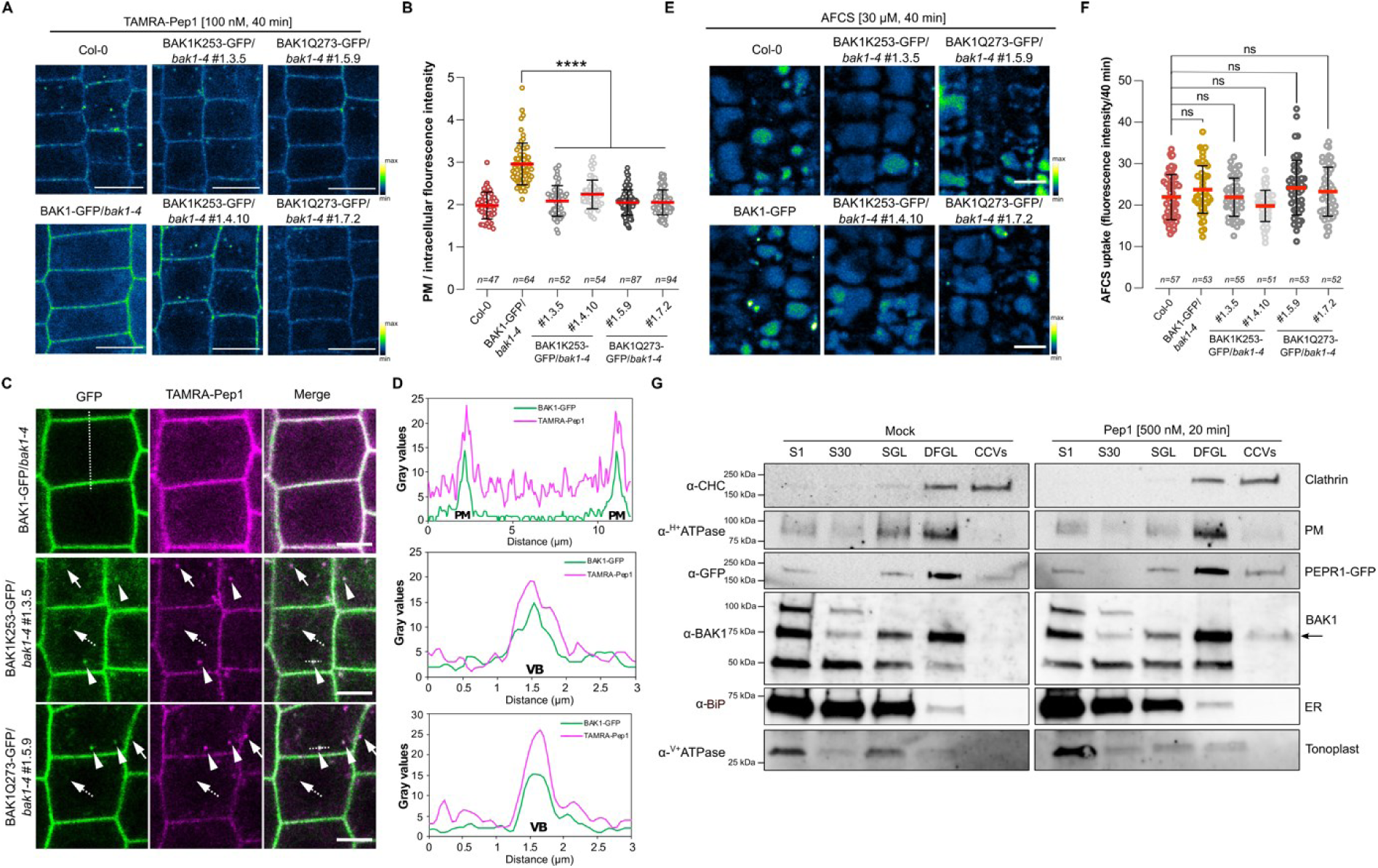
BAK1 and PEPR1 are likely co-internalized. **(A, E)** Representative images of TAMRA-Pep1 **(A)** and AFCS **(E)** root uptake visualized by color-based fluorescence intensity coding. Five-day-old seedlings of wild-type Col-0, *bak1-4*, *pBAK1-BAK1-GFP*/*bak1-4*, *pBAK1-BAK1K253-GFP*/*bak1-4* and *pBAK1-BAK1Q273-GFP/bak1-4* were treated with 100 nM TAMRA-Pep1 for 10 s (pulse) and with 40 µM of AFCS for 20 min (pulse), respectively, and imaged after 40 min (chase). Epidermal cells of root meristematic zone were imaged. Scale bars, 10 µm. **(B, F)** Fluorescence intensity measurements of the plasma membrane (PM) vs. intracellular compartments of the TAMRA-Pep1 uptake in **(A)** and AFCS uptake **(E)** after 40 min chase. Dots, red lines, and whiskers represent individual measurements, averages, and SD. The one-way ANOVA followed by Tukey’s test was used to determine significant differences. \*\*\*\**P*<0.0001; ns, not significant; *n*, number of cell analyzed. **(C)** Co-localization of BAK1-GFP and TAMRA-Pep1. Five-day-old *pBAK1-BAK1-GFP*/*bak1-4*, *pBAK1-BAK1K253-GFP*/*bak1-4* and *pBAK1-BAK1Q273-GFP/bak1-4* seedlings were treated with 100 nM of TAMRA-Pep1 for 10 s (pulse) and imaged after 40 min (chase). Arrowheads, arrows and dashed-arrows indicate co-localizing puncta, TAMRA-Pep1 puncta only and BAK1-GFP puncta only, respectively. Scale bars, 10 µm. **(D)** Line plot of the BAK1-GFP and TAMRA-Pep1 fluorescent signals of the dashed line in **(C)** VB, vesicle body; PM, plasma membrane. Epidermal cells of root meristematic zone were imaged. **(G)** Co-localization between PEPR1 and BAK1 in clathrin-coated vesicles (CCVs). CCVs were prepared from total plant extracts of 10-day-old *pRPS5A-PEPR1-GFP*/*pepr1pepr2* seedlings. Samples were collected during CCV purification and subjected to Western Blotting (WB) with antibodies against clathrin heavy chain (CHC), GFP, BAK1, and various subcellular organelle marker proteins. S1, supernatant after centrifugation at 1000*g*; S30, supernatant after centrifugation at 30,000*g*; SGL, sucrose step gradient load; DFGL, deuterium oxide/ficoll gradient load; CCVs, CCV-containing fraction. The following antibodies were used as organelle- or compartment-specific markers: *α*-H-ATPase (PM), *α*-V-ATPase (vacuole), and *α*-BiP (ER; endoplasmic reticulum).

To confirm that BAK1 and PEPR1 are internalized together, we purified clathrin-coated vesicles (CCVs) from Arabidopsis *pRPS5A-PEPR1-GFP*/*pepr1pepr2* seedlings and examined whether BAK1 was present alongside PEPR1 in the enriched CCV fractions after treatment with Pep1 (500 nM for 20 min). Whole Arabidopsis seedlings were used for the isolation of CCVs because Pep1 induced triggered PEPR1-GFP endocytosis in both shoots and roots (Supplemental Figure 9). Analysis of the enriched CCV fractions by immunoblotting revealed that BAK1 was barely detectable in the mock treatment but was enriched in CCVs after Pep1 treatment, as was PEPR1 (Figure 7G). Taken together, our data indicate that BAK1 and PEPR1 are likely endocytosed together.

## DISCUSSION

Endocytosis controls the amount of functional cell surface receptors and their ligands, acting as a spatial and temporal organizer of signaling (Sigismund et al., 2012). A link between endocytosis and signaling has been established for several Arabidopsis RKs involved in development and immunity (Erwig et al., 2017; Irani et al,. 2012; Mbengue et al., 2016; Ortiz-Morea et al., 2016). BAK1 and other SERK family members function as co-receptors (Fan et al., 2016), but experimental evidence for their involvement in the endocytosis of the main receptors is still lacking.

Transient expression analyses of BAK1 subcellular dynamics in cowpea mesophyll protoplasts (Russinova et al., 2004; Song et al., 2009) revealed that the C-terminal GFP-tagged BAK1 cycles between the PM and the endosomes. Surprisingly, similar dynamics was not observed in Arabidopsis roots (Bücherl et al., 2013), although, BAK1-GFP localized in the PM, and was able to complement the BR-related defects of the *bak1-4* null mutant (Ntoukakis et al., 2011; Lozano-Durán et al., 2013). Further assessment of BAK1 subcellular dynamics has been hampered by the observation that the C-terminal GFP-tagged BAK1 is unable to respond to exogenous flg22 and elf18 (Ntoukakis et al., 2011; Lozano-Durán et al., 2013). Here, we show that BAK1-GFP is also impaired in Pep1-triggered responses. Disruption of protein functionality due to a tag fusion has been previously reported in both plant and mammalian systems (Wiśniewska et al., 2006; Hammond et al., 2010; Hurst et al., 2018) and can potentially lead to misinterpretation of the results. Moreover, the functionality of protein fusions may depend on the protein complexes and their respective functions. One possible explanation for the impaired immune responses caused by the C-terminal GFP tag of BAK1 is that it may act in a dominant-negative manner, thereby interfering with receptor complex activation or downstream signal transduction. For example, it could disrupt the recruitment of certain protein partners to the receptor complex, affect PTMs crucial for signaling, or, similar to BAK1-5 protein, stabilize the PEPR1-BAK1 complex at the PM (Schwessinger et al., 2011). Consistent with this, the C-terminal GFP tag was found to impede the phosphorylation on S612 in the C-terminal tail of BAK1, the domain specifically targeted by phosphorylation events required to trigger immune responses in Arabidopsis in response to flg22 and elf18 (Perraki et al., 2018). Additionally, BAK1^S612A^ appeared to co-purify with more FLS2 than the wild-type BAK1 (Perraki et al., 2018). Remarkably, Pep1-induced PEPR1 endocytosis was also severely impaired in the transgenic lines expressing the C-terminal GFP-tagged BAK1, whereas the BRI1-dependent internalization of AFCS remained unaffected. Furthermore, BAK1^AAA^ and BAK1^S612A^ mutants deficient in conserved phosphorylation sites, significantly reduced Pep1-induced PEPR1 internalization, providing evidence that BAK1 phosphorylation, necessary for the immune responses (Perraki et al., 2018), is also important for Pep1-triggered PEPR1 endocytosis. Supporting this, the semi-dominant *bak1-5* mutant, known to disrupt the immune phosphorylation-dependent function of BAK1 (Perraki et al., 2018), strongly blocked Pep1-triggered PEPR1 endocytosis. Moreover, PEPR1 internalization was only slightly disturbed in the *bak1-4* null mutant, but severely affected in the *bak1-4serk1-1* mutants, suggesting that BAK1 and SERK1 redundantly regulate PEPR1 endocytosis, consistent with earlier studies in tobacco (Mbengue et al., 2016).

Examination of the co-localization between the C-terminal GFP-tagged BAK1 and TAMRA-Pep1 revealed that BAK1 localized to the PM, but it did not label the positive compartments of TAMRA-Pep1. This suggest that BAK1 dynamics may be a prerequisite for PEPR1, but not BRI1, endocytosis and signaling, consistent with observations that the C-terminal GFP-tagged BAK1 was functional in terms of BR responses (Ntoukakis et al., 2011; Lozano-Durán et al., 2013). To support this hypothesis and overcome the lack of immune responses caused by the C-terminal GFP tag, we applied an insertional GFP approach to identify a functional fluorescently tagged BAK1 for subcellular dynamics studies. Since the C-terminal tail of BAK1 is important for immune signaling (Wu et al., 2018; Perraki et al., 2018), we inserted the GFP before the critical residues within the C-terminal tail for the generation of BAK1M580-GFP and BAK1A593-GFP, and between the JMD and the KD for that of BAK1K253-GFP and BAK1Q273-GFP.

Remarkably, both BAK1K253-GFP and BAK1Q273-GFP versions retained partial responses to exogenous BRs and Pep1, showed dynamics, and restored PEPR1 endocytosis. In addition, BAK1K253-GFP and BAK1Q273-GFP co-localized with TAMRA-Pep1 puncta, suggesting that PEPR1 is endocytosed together with BAK1. Consistent with this co-localization, BAK1 and PEPR1 were detected together in isolated CCVs from Arabidopsis plants after Pep1 treatment. The fact that BAK1 is endocytosed together with PEPR1 raises the possibility that the BAK1-PEPR1 receptor complex maintains signaling from endosomes (Figure 8). In mammals, the role of endosomes as signaling platforms is a well-studied, albeit still controversial, phenomenon (Pavlos and Friedman, 2017). However, conclusive evidence for endosomal signaling in plants is still lacking. Further study of the internalization and dynamics of the PEPR1-BAK1 receptor complex may provide additional proof for the existence of similar mechanisms in plants.

**Figure 8.**
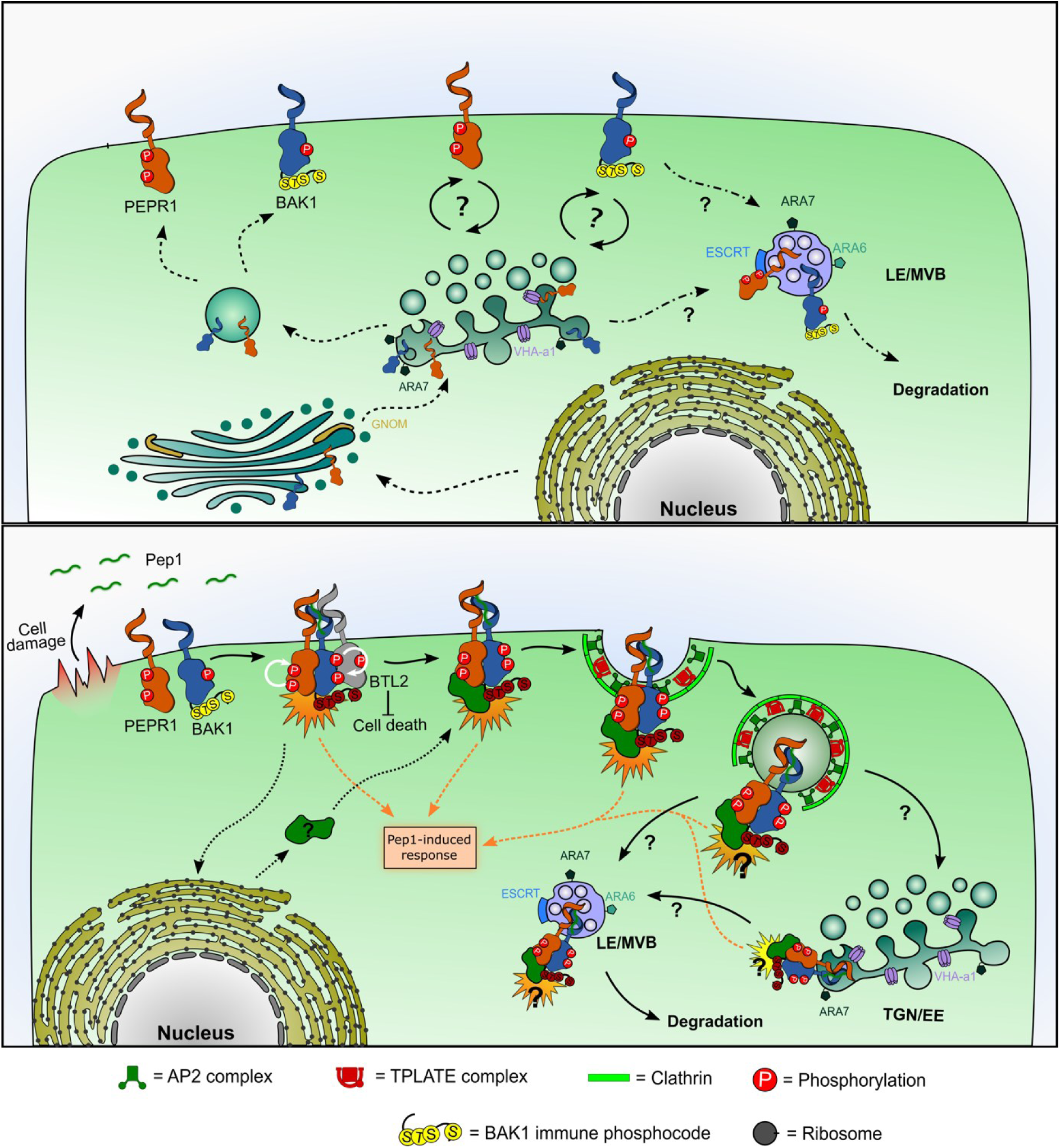
Proposed model of BAK1-PEPR1 activation and endocytosis. (A) In the non-elicited state, BAK1 and PEPR1 are delivered to the plasma membrane (PM) via the secretory pathway (dashed lines), passing through the endoplasmic reticulum (ER), Golgi apparatus, and trans-Golgi network/early endosomes (TGN/EE). Once at the PM, BAK1 and PEPR1 may undergo recycling. However, neither receptor is abundant in brefeldin A (BFA) bodies, suggesting that, if recycling occurs, it proceeds through a BFA-insensitive route. Ultimately, BAK1 and PEPR1 can be degraded (dash-dot lines) either by direct trafficking from the PM to late endosomes/multivesicular bodies (LE/MVBs) or via the TGN/EE and then to LE/MVBs for delivery to the vacuole, where they are degraded. (B) Upon Pep1 perception, PEPR1 dimerizes with BAK1 and trans-phosphorylates it. This presumably triggers phosphorylation of BAK1 at residues S602, T603, S604, and S612, which are required for Pep1-induced immune responses and for PEPR1 endocytosis. In parallel, BAK1 promotes increased PEPR1 phosphorylation, whereas PEPR1 subsequently attenuates BAK1 activity. Concurrently, BAK1 phosphorylates BAK-TO-LIFE 2 (BTL2), thereby suppressing BTL2-mediated autoimmunity. Pep1 signaling also induces the transcription and translation of an unknown factor that may be required for endocytosis of the PEPR1–Pep1–BAK1 complex. PEPR1 is internalized together with BAK1 via clathrin-mediated endocytosis. After internalization, the complex is targeted directly to LE/MVBs or traffics via the TGN/EE and then to LE/MVBs for delivery to the vacuole for degradation.

The ligand-induced endocytosis of PEPR1 depended on its kinase activity, but the phosphorylation relationship between BAK1 and PEPR1 is unclear. Protein kinases lacking Arginine in the catalytic loop HxD motif, also known as non-RD kinases, are implicated in immune signaling and may be activated by phosphorylation-dependent conformational changes in the receptor cytoplasmic domain, but not by an activation loop phosphorylation (Bender et al., 2021). For example, BAK1 phosphorylates the non-RD kinase EFR in the activation loop to stabilize its active conformation, allowing EFR in turn to allosterically activate BAK1 but not vice versa (Wang et al., 2014; Schwessinger et al., 2011; Mühlenbeck et al., 2024). Unlike other PRRs, PEPR1 is an RD kinase with strong kinase activity *in vitro* and has been shown to phosphorylate various partners, such as BIK1 (Liu et al., 2013) and the REGULATOR OF G SIGNALLING PROTEIN1 (RGS1) (Tunc-Ozdemir et al., 2017). BAK1 can trans-phosphorylate most of its co-receptors, including the non-RD kinases FLS2 (Lin et al., 2014), EFR (Wang et al., 2014; Schwessinger et al., 2011), the RD kinase BRI1 (Wang et al., 2008), and other substrates, such as NUCLEAR SHUTTLE PROTEIN-INTERACTING KINASE1 (NIK1) (Li et al., 2019), BIK1 (Liu et al., 2017), and H^+^-ATPASE ISOFORM2 (AHA2) (Pei et al., 2022). We found that PEPR1 can phosphorylate BAK1 *in vitro*, but, surprisingly, BAK1 did not phosphorylate PEPR1 *in vitro* and *in vivo* (Supplemental Figure 3B; Supplemental Dataset 2). However, the presence of the BAK1 kinase visibly increased PEPR1 phosphorylation *in vitro*, comparable to the observed *in vivo* increase in phosphorylation at T896 in PEPR1 when co-expressed with BAK1 in the presence of Pep1 in tobacco (Supplemental Dataset 2). Additionally, PEPR1 kinase reduced BAK1 phosphorylation *in vitro*. Likewise, FLS2 and EFR reduced BAK1 kinase activity *in vitro* (Wang et al., 2014). Taken together, the mutual trans-phosphorylation mechanism for activation of RKs complexes, as in BRI1-BAK1 and FLS2-BAK1, might not apply to PEPR1-BAK1 and requires further investigation.

In conclusion, our study demonstrates that the PEPR1 endocytosis is controlled by the co-receptor BAK1 via phosphorylation in its C-terminal tail, which is indispensable for Pep1-mediated signaling. This raises the question of the possible mechanism underlying this regulation (Figure 8). It is tempting to speculate that BAK1 counteracts an as-yet-unknown component of the receptor complex that negatively regulates PEPR1 endocytosis. In line with this hypothesis, BAK1 restricts the activation of the LRR RK, BAK-TO-LIFE 2 (BTL2) through specific phosphorylation to maintain cellular integrity (Yu et al., 2023). However, whether BTL2 affects endocytosis of the main receptor remains unclear. BAK1 can also regulate specific PTMs in PEPR1 that positively regulate ligand-induced receptor endocytosis, such as phosphorylation or ubiquitination (Dubeaux and Vert, 2017). Notably, in contrast to other BAK1 receptor partners, PEPR1 can phosphorylate BAK1 to reduce its activity.

Additionally, the immune-required phosphorylation of BAK1 is not necessary for BRI1 endocytosis, suggesting that BAK1 employs different mechanisms to regulate different receptor partners. Nevertheless, the precise mechanism by which BAK1 controls PEPR1 endocytosis remains to be elucidated. Whether BAK1 has a similar effect on other PRRs is unclear. Some conflicting evidence exists that BAK1 is essential for ligand-induced endocytosis of certain PRRs in transient assays (Mbengue et al., 2016), but it is not required for FLS2 recycling in Arabidopsis leaves, regardless of the ligand presence (Beck et al., 2012). Furthermore, using the newly designed and partially functional BAK1-GFP lines, it would be interesting to investigate BAK1 dynamics within different immune receptor complexes, such as FLS2 and EFR. Our findings suggest that the ligand-induced endocytosis of the main receptor is regulated in a co-receptor-dependent manner.

## METHODS

### Plant material, growth conditions and reagents

All mutants and transgenic lines used in this study were in the background of the *Arabidopsis thaliana* (L.) Heynh. (accession Columbia-0 [Col-0]). Seeds were surface-sterilized with chlorine gas and then placed on plates with half-strength Murashige and Skoog (½MS) medium containing 0.8% (w/v) agar, and 2.5 mM methyl ester sulfonate at pH 5.7. After vernalization for 2 days at 4°C, the plates were moved to the growth chamber at 22°C in a 16 h/8 h light/dark cycle with a light intensity of 120 μE m^-2^ s^-1^. The following mutant and transgenic Arabidopsis lines have been described previously: *pRPS5A-PEPR1-GFP*/*pepr1pepr2* (Ortiz-Morea *et al*, 2016); *bak1-4* (Chinchilla *et al*, 2007); *bak1-5* (Schwessinger *et al*, 2011); *serk1-1bak1-4* (Meng *et al*, 2015); *bak1-3/serk4-1* (Du *et al,* 2016); *serk1-1(+/-)serk2-1bak1-4* (Meng *et al*, 2015); *serk1-1serk2-1(+/-)serk4-1* (Meng *et al*, 2015); *pBAK1-BAK1*/*bak1-4* (Perraki *et al*, 2018); *pBAK1-BAK1^S612A^*/*bak1-4* (Perraki *et al*, 2018); and *pBAK1-BAK1^S602A;T603A;S604A^*/*bak1-4* (Perraki *et al*, 2018). The *pRPS5A-PEPR1-GFP* construct (Ortiz-Morea *et al*, 2016) was introduced into the *bak1-4* mutant (Chinchilla *et al*, 2007) by Agrobacterium-mediated transformation. The *pRPS5A-PEPR1-GFP*/*pBAK1-BAK1^S612A^*/*bak1-4* and *pRPS5A-PEPR1-GFP*/*pBAK1-BAK1^S602A;T603A;S604A^*/*bak1-4* lines were generated by crossing *pRPS5A-PEPR1-GFP/bak1-4* with either *pBAK1-BAK1^S612A^*/*bak1-4* or *pBAK1-BAK1^S602A;T603A;S604A^*/*bak1-4*. For phenotypic analysis, plants were grown in soil in a growth chamber at 22°C, 58% relative humidity, and a 16 h light/8 h light/dark photoperiod for 6 weeks. N-(3-triethylammoniumpropyl)-4-(6-(4-(diethylamino) phenyl) hexatrienyl) pyridinium dibromide (FM4-64; 2 mM stock in purified water; Invitrogen), BFA [50 mM stock in dimethyl sulfoxide (DMSO); Sigma-Aldrich], CHX (50 mM stock in DMSO; Sigma-Aldrich), BL (10 mM stock in DMSO; Wako Pure Chemical Industries and OlChemlm) were used at the concentrations indicated in the figure legends.

### Plasmid construction

cDNA or genomic DNA from Arabidopsis Col-0 was used as template for PCR cloning. The *BAK1* promoter was amplified and introduced into the *pDONRP4-P1R* donor vector to build *pDONRP4-P1R-pBAK1*. The *BAK1* coding sequence without stop codon was cloned into *pDONR221* to generate *pDONR221-BAK1* that was used as a template together with *pDONRP2-P3R-GFP* to generate the BAK1 insertional GFP versions by overlapping PCR and subcloned into *pDONR221* to generate *pDONRP1P2-BAK1K253-GFP, pDONRP1P2-BAK1Q273-GFP, pDONRP1P2-BAK1M580-GFP* and *pDONRP1P2-BAK1A593-GFP.* The destination vectors were generated by recombining the *pH7m24GW,3, pDONRP4-P1R-pBAK1,* and *pDONRP1P2-BAK1-GFP* versions.

For protein purification, the CD-encoding sequences of BAK1were amplified and introduced into *pDONR221* to generate *pDONR221-BAK1-CD* that was used as template to generate *BAK1-Km(D416N)-CD* by overlapping PCR and subcloned into *pDONR221*. The entry clones *pDONR221-BAK1-CD* and *pDONR221-BAK1-Km-CD* were recombined into the bacterial expression vector *pOPINM* (Berrow *et al*, 2007) containing the His6-MBP tag. Sequences encoding CDs of *PEPR1* and *PEPR1-Km(K855E)* were amplified with specific primers containing restriction sites. The point mutations were generated by means of overlapping PCR. The PCR products and the plasmid *pGEX-5×1* were digested with the restriction enzymes EcoRI and BamHI (FastDigest; Thermo Fisher Scientific). The linearized and purified plasmid was dephosphorylated by Thermosensitive Alkaline Phosphatase (Promega). PCR products and plasmid were ligated by T4 DNA ligase (Invitrogen) to generate *pGEX-5×1-PEPR1-CD* and *pGEX-5×1-PEPR1-Km-CD*. Constructs were introduced into the *Escherichia coli* strain Rosetta2 for protein production. All primers used are listed in Table EV1. All clones were confirmed by sequencing.

### Generation of transgenic plants

Plants were transformed with the *Agrobacterium tumefaciens* strain C58C1 by means of the floral dip method (Clough & Bent, 1998). BAK1 constructs were transformed into the *bak1-4* null mutant Transformed plants were selected on ½MS agar medium containing 50 mg/l hygromycin.

### Peptides

The peptide Pep1 (ATKVKAKQRGKEKVSSGRPGQHN) with a HPLC purity of 95% and the peptide Pep1 labelled with 5’-carboxytetramethylrhodamine at the N-terminal (TAMRA-Pep1) with a HPLC purity of 95% were purchased from EZBiolab. The peptides were dissolved in water to obtain peptide stocks of 1 mM and further diluted with ½MS medium.

### Root and hypocotyl growth assays

Seeds were sown on ½MS solid medium, stratified for 2 days at 4°C in the dark, and placed vertically in the light. Four days after germination, seedlings were transferred to solid ½MS medium supplemented with or without the indicated amount of Pep1 and incubated for a further 3 days, whereafter the plates were scanned. For the hypocotyl, seeds were sown directly on ½ MS medium supplemented with either increasing concentrations of BL or (DMSO), stratified for 2 days at 4°C in the dark. The plates were moved to growth chamber at 22°C and placed vertically in light for 6 h. Then, the plates were wrapped in aluminum foil for dark condition and kept vertically in growth chamber at 22°C for 5 days. For root and hypocotyl measurements, plates were scanned and the scanned images were processed and evaluated with the ImageJ software and plotted relative to the untreated control.

### Detection of reactive oxygen species (ROS) burst

For ROS burst assay (Zhang *et al*, 2010), leaf discs (5 mm in diameter) were collected from the third or fourth true leaves of four-week-old Arabidopsis plants grown in soil. These discs were placed into individual wells of a 96-well plate, each containing 100 μL of distilled water, and left to incubate overnight under a 12-hour light/12-hour dark cycle. The next day, the water was replaced with 100 μL of a reaction solution composed of 50 μM luminol and 10 μg/mL horseradish peroxidase (Sigma-Aldrich, USA), with or without the addition of Pep1 at concentrations as indicated. Luminescence signals were detected using a GloMax-Multi System luminometer (Promega, USA).

### MAP kinase assay

Arabidopsis seedlings were germinated on ½ MS medium and, after three or five days, were transferred to a 12-well plate with ½ MS medium. Following 1 hour of conditioning, the medium was supplemented with 100 nM Pep1. Seedlings were incubated with peptides for the indicated time points, whereafter they were harvested and frozen in liquid N_2_. Frozen plant material was ground and the extraction buffer [50 mM Tris, pH 7.5, 200 mM NaCl, 1 mM EDTA, 10% (v/v) glycerol, 0.1% (v/v) Tween 20, 1 mM phenylmethylsulfonyl fluoride, 1 mM DTT, 1×PhosSTOP™ (4906837001; Sigma-Aldrich) and 1×cOmplete™ ULTRA Tablets (5892791001; Sigma-Aldrich)] was added at the ratio of 2 µL per mg tissue (1:2 w:v). Samples were vortexed and centrifuged at 13000*g* for 30 min. Protein concentration was quantified in the supernatant with the Qubit protein assay kit (Thermo Fischer Scientific). Equal amounts of proteins (30 μg) were separated on 7.5% sodium dodecyl sulphate (SDS)/polyacrylamide gel electrophoresis (PAGE) and transferred onto a nitrocellulose membrane (Bio-Rad). After blocking with 3% bovine serum albumin, membranes were incubated overnight with primary antibodies against anti-phospho-p44/42 (1:2500; Cell Signaling Technology). Horseradish peroxide (HRP)-conjugated anti-rabbit secondary antibody (1:10000; GE Healthcare;) was used and signals were detected with the SuperSignal™ West Femto Maximum Sensitivity Substrate (34095; Thermo Fisher Scientific). Antibodies were stripped in 1:1 (v:v) 10% SDS and 100 mM glycine-HCl (pH 2.5) solution, washed five times, and reblotted with anti-MPK6 antibody (1:8000; Sigma-Aldrich) and anti-MPK3 antibody (1:2500; Sigma-Aldrich). HRP-conjugated anti-rabbit secondary antibody (1:10000; GE Healthcare) was used and signals detected with Western Lightning Plus Electrochemiluminescence (NEL105001EA; PerkinElmer). All blots were imaged in a ChemiDoc XRS+ imaging system (Bio-Rad).

### Immunoblots

The Arabidopsis seedlings were ground in liquid N_2_, resuspended in total protein extraction buffer (25 mM Tris-HCl, pH 7.5, 150 mM NaCl, and Roche cOmplete ULTRA protease inhibitor cocktail, Roche PhosSTO tablet) in a 1:2 (w/v) ratio, and centrifuged at 15,000*g*. The supernatants were mixed with required amount of 4× NuPAGE lithium dodecyl sulphate (LDS) sample buffer (Invitrogen) and 10× NuPAGE sample-reducing agent (Invitrogen), heated at 70°C for 10 min, and loaded onto 4-20% Mini-PROTEAN TGX precast gels. The proteins were transferred to nitrocellulose membranes by means of the Trans-Blot® Turbo™ Transfer System (Bio-Rad). For protein detection, the following antibodies were used: monoclonal α-GFP-HRP (1/5,000; Miltenyi Biotech), monoclonal α-tubulin (1/10,000; Sigma-Aldrich), α-BES1 (1:4,000; Yin et al., 2002), α-BAK1 (1/5,000; custom-made by Eurogentec). Blots were developed with Western Lightning Plus-ECL, Enhanced Chemiluminescence Substrate (Perkin-Elmer), and imaged with the Bio-Rad ChemiDoc XRS+molecular imager.

### TAMRA-Pep1 internalization assay and image analysis

Five-day-old seedlings were dipped into 500 μl of 100 nM TAMRA-Pep1 dissolved in ½MS medium for 10 sec, washed with ½MS liquid medium three times, and kept in 500 μl of ½MS medium over a Parafilm piece in a Petri plate. The epidermal cells of the root meristematic zone were imaged at the indicated time points with a laser scanning confocal microscope SP8X (Leica) equipped with an HC PL APO CS2 40×/1.10 water-corrected objective with 4× digital zoom. The excitation wavelength was 559 nm by white light laser. Emission was detected at 570-670 nm by hybrid detectors (Leica). For the image quantification with FIJI (https://imagej.net/software/fiji/), the first images were converted to 8-bit images; subsequently, the entire plasma membranes of individual cells were selected with the brush tool size 10 pixels and the intracellular spaces with the polygon selection tool. The number of the 5% highest pixels was obtained for both PM and intracellular space. These values were used to obtain a ratio. Ten epidermal cells from five plants were quantified.

### AFCS uptake assays and image analysis

The AFCS uptake assay was done as previously described (Irani *et al*, 2014) with modifications. Six-day-old seedlings grown on solid ½MS were transferred to 200 µl of liquid ½MS on a Parafilm piece placed in a Petri plate, humidified with wet laboratory wipes for 10 min. The medium was replaced with ½MS supplemented with 30 µM AFCS for 40 min (pulse). Seedlings were washed six times and chased for 30 min on ½MS followed by imaging. Epidermal cells of the root meristematic zone were imaged with the laser scanning confocal microscope SP8X (Leica) equipped with an HC PL APO CS2 63×/1.20 water-corrected objective with 4× digital zoom. The excitation wavelength was 633 nm by white light laser. Emission was detected at 650-700 nm by hybrid detectors (Leica). For fluorescence signal measurements in vacuoles, stacks of four to six slices of 1.5 μm were obtained. Images were analyzed and signals quantified with the Fiji/ImageJ software. Four to six slices covering the epidermal cell layer were merged by means of maximum projection. The images were processed with Gaussian blur filter (1) and Subtraction Background (100). Rectangular regions of interest were determined omitting over-stained regions, such as the lateral root cap. The signal intensity was quantified in seven to nine seedlings per genotype.

### Protein production and purification

For GST-tagged protein production, the *E. coli* strain Rosetta2 harboring each of the plasmids pGEX-5×1-PEPR1-CD, pGEX-5×1-PEPR1-Km-CD or *pDEST15-BIK1-Km(KK105AA*) was grown in Luria-Bertani (LB) medium supplemented with carbenicillin (100 µg/ml) at 37°C until an optical density OD_600_ of 0.6-0.8. Protein production was induced with 1 mM isopropyl β-D-1-thiogalactopyranoside (IPTG) at 16°C overnight, and shaking at 200 rpm. The bacterial culture was pelleted (15 min, 5000*g*) and frozen at −20°C. The cells were resuspended in extraction buffer [50 mM Tris-HCl, pH 8.5, 100 mM NaCl, 1 mM EDTA, 1 mM DTT, protease inhibitor cocktail tablet (cOmplete ULTRA Tablets; Roche), 1 mg/ml lysozyme, 0.4 mM PMSF, 1% Triton X-100] and incubated on ice for 60 min. Resuspended cells were sonicated until the solution became fluid and clear. The lysate was centrifuged (30 min, 12000*g* at 4°C) and the supernatant was transferred to a new tube. Proteins were purified with Glutathione Sepharose^®^ 4B (GE Healthcare). Glutathione Sepharose slurry (300 µl) was washed twice with binding buffer (50 mM Tris-HCl, pH 8.5, 100 mM NaCl, 1 mM EDTA, and 1 mM DTT) by addition of 5 ml buffer for 1 ml slurry. After washing, the slurry was added to the lysate and incubated for 60 min on an end-to-end rotor at 4°C. After incubation, slurry was spun down (5 min, 500*g*) and washed twice with 10 ml washing buffer A (50 mM Tris-HCl, pH 8.5, 100 mM NaCl, 1 mM EDTA, 1 mM DTT, and 0.1% [v/v] Triton X-100) for 15 min on an end-to-end rotor at 4°C. The slurry was further washed twice with 10 ml binding buffer (50 mM Tris-HCl, pH 8.5, 100 mM NaCl, 1 mM EDTA, 1 mM DTT) for 15 min on an end-to-end rotor at 4°C. For elution, the slurry was incubated for 10 min at room temperature on an end-to-end rotor with 500 µl elution buffer (50 mM Tris-HCl, pH 8.5, 100 mM NaCl, 1 mM EDTA, 1 mM DTT, 20 mM freshly prepared gluthatione, pH 8.0). The elution was repeated three times and the three eluates were pooled and concentrated with Amicon^®^ Ultra 30K (Millipore) until 500 µl was obtained.

For His6-MBP-tagged proteins, *E. coli* strain Rosetta2 harboring the plasmid *pOPNIM-BAK1-CD* and *pOPNIM-BAK1-Km-CD* were grown in LB medium supplemented with 100 µg/ml carbenicillin at 37°C until OD_600_ = 0.6-0.8. Protein production was induced with 1 mM of IPTG at 16°C overnight, with shaking at 200 rpm. The bacterial culture was pelleted (15 min, 5000*g*) and frozen at −20°C. The cells were resuspended in extraction buffer [20 mM Tris-HCl, pH 8.5, 200 mM NaCl, 1 mM EDTA, 1 mM DTT, protease inhibitor cocktail tablet (cOmplete ULTRA Tablets; Roche), 1 mg/ml lysozyme, 0.4 mM PMSF] and incubated on ice for 60 min. Resuspended cells were sonicated until the solution became fluid and clear. The lysate was centrifuged (30 min, 12000*g* at 4°C) and the supernatant was transferred to a new tube. Proteins were purified with mylose resin (NEB). Amylose slurry was washed three times with wash buffer (20 mM Tris-HCl pH 8.5, 200 mM NaCl, 1 mM EDTA, and 1 mM DTT) and then incubated with the lysate for 60 min on an end-to-end rotor at 4°C. After incubation, the slurry was spun down (5 min, 500*g*) and washed three times with 10 ml of wash buffer for 15 min on an end-to-end rotor at 4°C. For elution, the amylose slurry was incubated for 10 min at room temperature on an end-to-end rotor with 1 ml elution buffer (20 mM Tris-HCl pH 8.5, 200 mM NaCl, 1 mM EDTA, 1 mM DTT, and 10 mM maltose). Elution was repeated three times and the three eluates were pooled and concentrated with Amicon^®^ Ultra 30K (Millipore) until 500 µl was obtained.

### *In vitro* kinase assay and in-gel trypsin digestion for mass spectrometry (MS)

The *in vitro* kinase assay was carried out by mixing either GST-PEPR1-CD or GST-PEPR1-Km-CD with His6-MBP-BAK1-CD; all the combinations, including the proteins alone, were tested for autophosphorylation. CDs were mixed with 5 µl of [γ-^32^P] ATP (1 µCi/µl) and the kinase assay buffer (50 mM Tris-HCl, pH 7.5, 100 mM NaCl, 10 mM MgCl_2_, and 10 µM ATP) was added in 30 µl final volume. The reaction was incubated for 1 h at 25°C and stopped by addition of 10 µl loading buffer 4× [NuPAGE® LDS Sample Buffer (4×)] and 5 µl of 10× reduction buffer [NuPAGE® Sample Reducing Agent (10×)]. Samples were loaded on 4-20% precast SDS-PAGE gels (Bio-Rad). The gels were stained with Bio-Safe Coomassie Brilliant Blue (CBB) (Bio-Rad), dried between cellophane plastic sheets, and [γ-^32^P] was detected by autoradiography. To examine the phosphorylation sites by MS, an *in vitro* kinase experiment with non-radioactive ATP was conducted after the previously described steps. After the kinase reaction had ended, the samples were segregated by SDS-PAGE and then stained with CBB. Thereafter, in-gel trypsin digestion was carried out, as outlined previously. The resulting digested peptides were purified with OMIX C18 Pipette Tips (Agilent Technologies) and subsequently subjected to a liquid chromatography-tandem MS (MS/MS).

### Detection of protein phosphorylation in the *Nicotiana benthamiana* leaf epidermis

Agrobacteria containing *P19*, *p35S-PEPR1-Km-GFP* or *p35S-BAK1-mCherry* were co-infiltrated into *Nicotiana benthamiana* leaves with the combination shown in the figure. After 60 hours, leaves were treated with water or 500 nM Pep1 for 15 min by injection before harvesting. Four biologically replicated samples per experimental group were ground with liquid nitrogen. Total protein was extracted from 5 mL of compacted plant powder using extraction buffer [50 mM Tris-HCl pH7.5, 150 mM NaCl, 1 mM EDTA, 1% NP-40, 5% glycerol, protease inhibitor cocktail (Roche) and PhosSTOP (Roche)]. Immunoprecipitation was performed with GFP-Trap magnetic agarose (ChromoTek) followed by 3 washes with wash buffer (50 mM Tris-HCl pH7.5, 150 mM NaCl, 1 mM EDTA and 1% NP-40) and 1 wash with 50 mM TEAB pH8.0 (Thermo Fischer). Protein was eluted on beads using 0.5 µg trypsin, and then alkylated of cysteine residues using 15 mM tris(carboxyethyl)phosphine (TCEP, Pierce) and 30 mM iodoacetamide (Sigma-Aldrich). The peptides were completely digested by incubation with 0.5 µg trypsin for 16 hours and desalting with OMIX C18 pipette tips (Agilent Technologies). The LC-MS/MS analyses were carried out as described previously (Vu *et al*. 2021). Peptide searches were performed using MaxQuant (version 2.1.4.0) on a high performance computer (HPC-UGent) against a database containing all *Nicotiana benthamiana* protein sequences from UniProt (Proteome ID: UP000084051) and the *Arabidopsis thaliana* protein sequences of PEPR1-Km-GFP and BAK1-mCherry.

### Database searching and data analysis

MS/MS spectra were searched with the BL21(DE3) proteome database and the sequences of GST-PEPR1-CD, GST-PEPR1-CD-kinase dead, HIS6-MBP-BAK1-CD, and HIS6-MBP-BAK1-Km-CD with the MaxQuant software (version 1.6.17.0) by means of the UseGalaxy.be server. MaxQuant settings can be found in Dataset EV1. The ‘Phospho (STY).txt’ output file generated by the MaxQuant search was loaded into the Perseus software (version 1.6.7.0) for analysis. All the data were selected first by a localization probe cut-off of >0.75. The Log2-transformed phosphosite intensities were used for further analysis. The ‘proteinGroups.txt’ file was also loaded into the Perseus software (version 1.6.7.0) for analysis. The label-free quantification (LFQ) intensities were loaded and transformed by Log2 for further analysis.

### CCV Purification

Seven-day-old *pRPS5A-PEPR1-GFP*/*pepr1pepr2* seedlings were grown in ½MS. A sample of 30 g was ground at 4°C and fractionated to purify CCVs as described previously (Mosesso *et al*, 2019). Samples collected during the purification and the final CCV fraction were analyzed by immunoblot analysis with antibodies against organelles and/or cellular compartments. The antibodies used were: HRP-coupled monoclonal α-GFP (1:2,500; Miltenyi Biotech), α-CHC (1:5,000; Santa Cruz Biotechnologies), α-H^+^ ATPase (1:5,000; Agrisera), α-V-ATPase (1:5,000; Agrisera), and α-BiP (1:2,000; Agrisera).

### RT-qPCR

Total RNA was extracted from approximately 100 mg of 5-day-old seedlings with the RNeasy kit (Qiagen). cDNA was synthesized from 1000 ng of RNA with the qScript cDNA Supermix (Quantabio). The resulting cDNA was diluted to 1:5 with nuclease-free water. RT-qPCRs were run with SYBR green I Master kit (Roche) on a LightCycler 480 (Roche). Expression was normalized to that of *ACTIN2* and *GAPDH* genes. The cycling conditions were as follows: preincubation at 95°C for 10 min; 45 amplification cycles at 95°C for 10 s, 60°C for 15 s, and 72°C for 15 s; melting curve at 95°C for 1 s, and 65°C for 1 s, followed by cooling at 40°C for 10 s. The integrity of the PCR amplicon was confirmed by melting curve analyses. The primers used are listed in Table EV1.

### Quantification and statistical analysis

All statistical analyses were carried out in GraphPad Prism 9. Statistically significant differences were determined by one-way or two-way Analysis of Variance (ANOVA) test.

### Data availability

No data were deposited in external repositories. Data used for quantifications as well as full Western blots can be found in the source data file. All material will be made available upon request to the corresponding author.

## SUPPLEMENTAL INFORMATION

Supplemental information is available online

## AUTHOR CONTRIBUTIONS

L.A.N.C., F.A.O.-.M. and E.R. initiated and designed the experiments. L.A.N.C. did most of the work. X. X., S.-L. Y. and I.D.S. identified the *in vitro* / *in vivo* phosphorylation sites. S. Y. did phenotypic analysis. I.Y. performed some MAPK and ROS assays. I. D.S., P.H., L.S., and E.R. provided supervision, funding and analyzed the data. L.A.N.C. and E.R. wrote the manuscript and all authors revised it.

## Supporting information

SDataset 1

SDataset2

## ACKNOWLEDGEMENTS

We thank Yanhai Yin for providing the α-BES1 antibody, Cyril Zipfel for published materials, Guilherme Perreira de Oliveira and Simone Di Rubbo for technical help, Daniel Scherer de Moura and Daniel V. Savatin for constructive discussions, and Martine De Cock for help in preparing the manuscript.

## Supplemental information

The supplemental file includes 9 supplemental figures and 1 supplemental table.

**Supplemental Figure 1.**
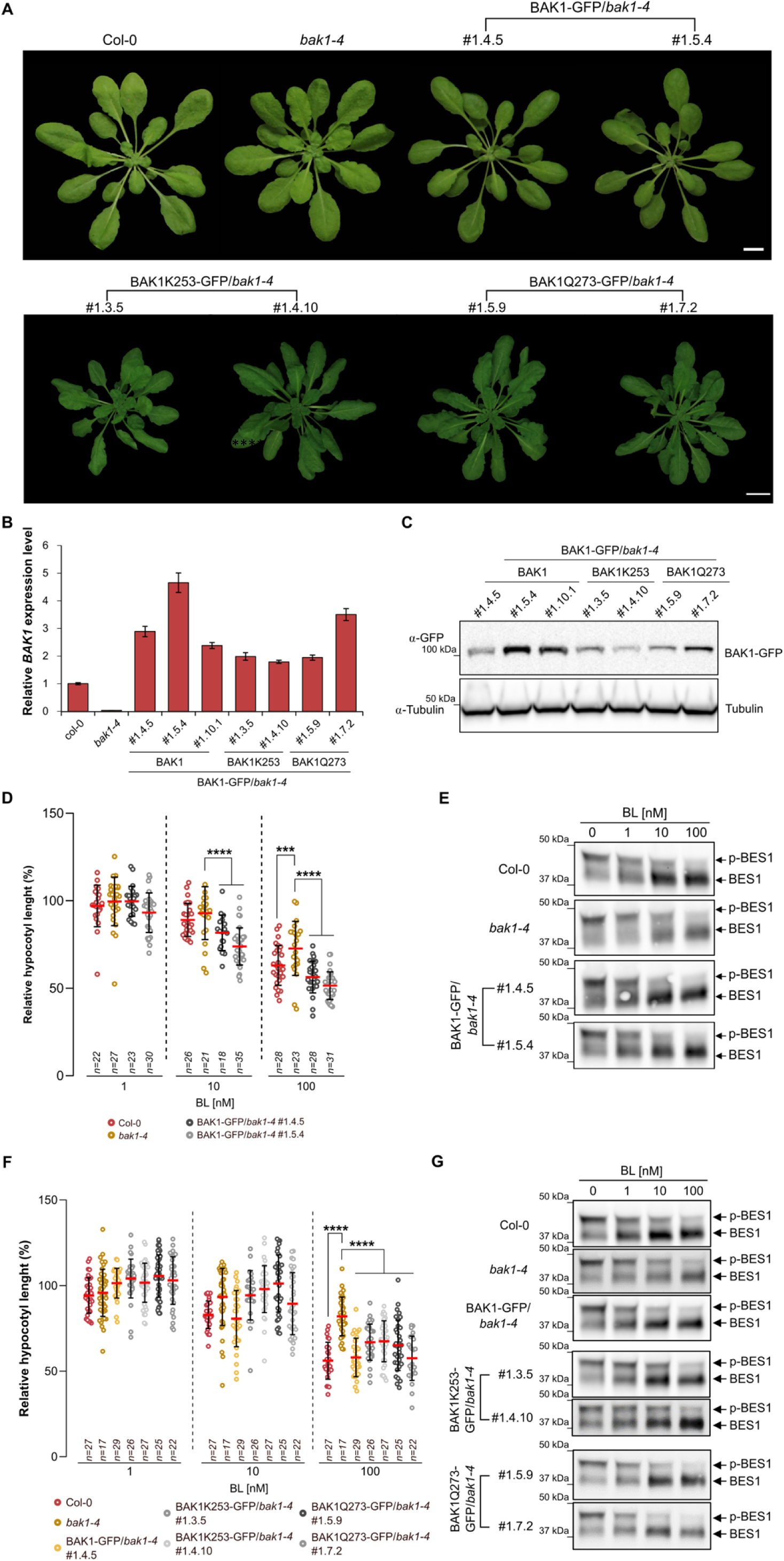
The C-terminal BAK1-GFP tag did not impair BR responses. **(A)** Representative images of 4-week-old plants of the indicated genotypes. Scale bars, 1 cm. **(B)** Real-time quantitative reverse transcription PCR (qRT-PCR) analysis of *BAK1*. Total RNA was isolated from 7-day-old seedlings of two or three independent transgenic lines. Transcript levels were normalized to *GAPDH* and *ACTIN2* gene expression and three biological with three technical replicates each were quantified. Data are means ± SD. **(C)** Total proteins isolated from 7-day-old seedlings and detected by Western Blotting (WB) with α-GFP antibodies to detect the BAK1-GFP. Numbers indicate individual transgenic lines. α-tubulin antibodies were used to equilibrate protein loading. **(D, F)** Hypocotyl length measurements after brassinolide (BL) treatment relative to the mock (DMSO). Seedlings of wild-type Col-0, *bak1-4* and two independent transgenic lines of each *pBAK1-BAK1-GFP*/*bak1-4*, *pBAK1-BAK1K253-GFP*/*bak1-4,* and *pBAK1-BAK1Q273-GFP*/*bak1-4* were grown for 4 days in the dark on ½MS agar plates supplemented with increasing concentrations of BL. Dots, red lines, and whiskers represent individual measurements, averages, and SD, respectively. Statistically significant difference were analyzed with two-way ANOVA followed by a Tukey’s test. \*\*\**P*<0.001; \*\*\*\**P*<0.0001; *n*, number of seedlings analyzed. **(E, G)** BES1 dephosphorylation assay after BL treatment. Five-day-old seedlings of wild-type Col-0, *bak1-4* and of two independent transgenic lines of each *pBAK1-BAK1-GFP*/*bak1-4*, *pBAK1-BAK1K253-GFP*/*bak1-4,* and *pBAK1-BAK1Q273-GFP*/*bak1-4* were treated with increasing concentrations of BL for 1 h. BES1 was detected by WB with the α-BES1 antibody and discrimination between phosphorylated (p-BES1) and unphosphorylated BES1 was based on the protein size shift.

**Supplemental Figure 2.**
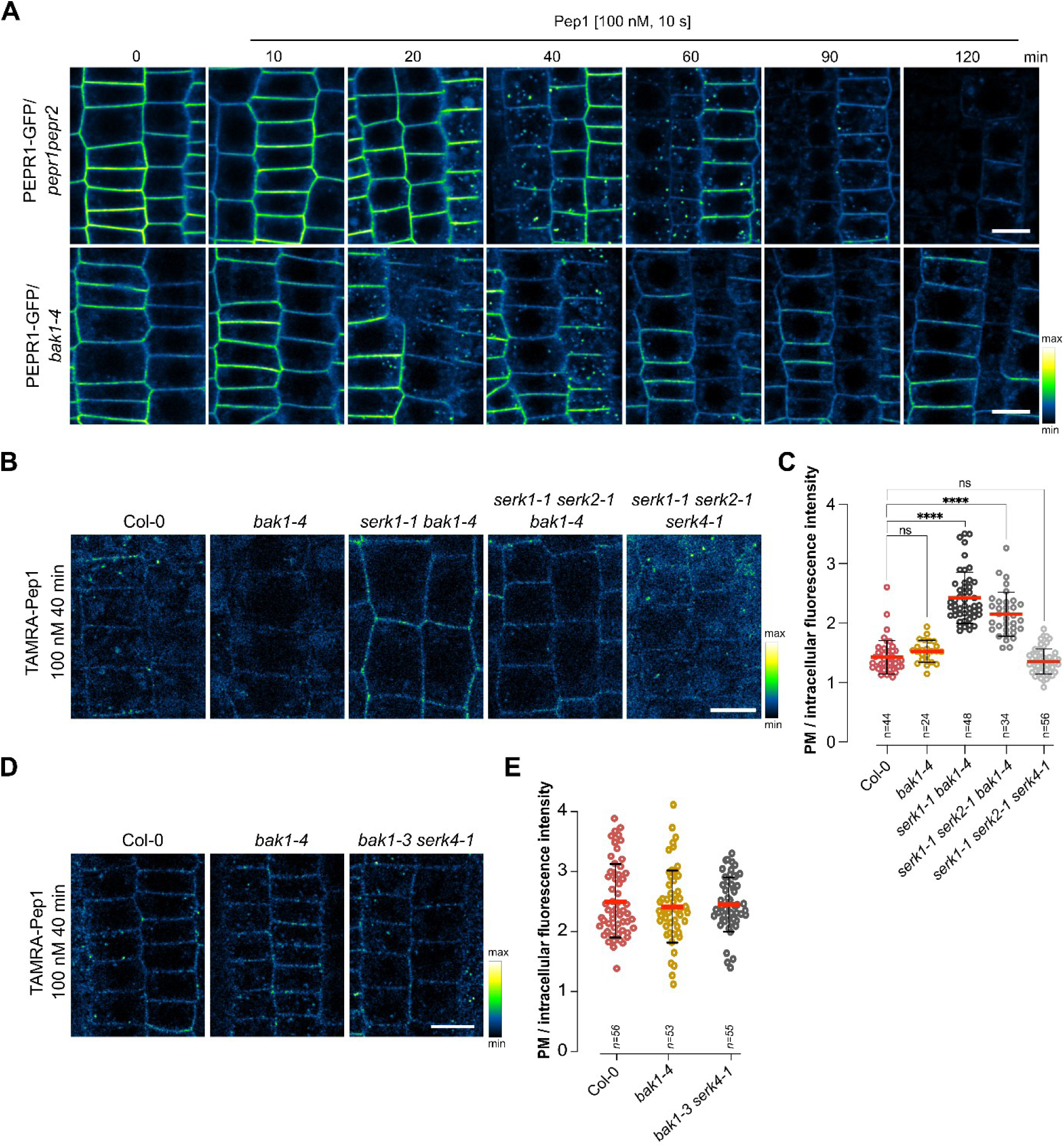
PEPR1 endocytosis requires SERKs. **(A)** Representative images of PEPR1-GFP endocytosis in roots of *pepr1pepr2* and *bak1-4* mutants. Five-day-old *pRPS5A-PEPR1-GFP/pepr1pepr2* and *pRPS5A-PEPR1-GFP*/*bak1-4* seedlings were treated with 100 nM Pep1 for 10 s (pulse) and imaged at the indicated time points (chase). (**B, D**) TAMRA-Pep1 root uptake visualized by color-based fluorescence intensity coding. 5-day-old wild-type Col-0*, bak1-4, serk1-1bak1-4*, *serk1-1serk2-1bak1-4* and *serk1-1serk2-1serk4-1* (**B**) and Col-0, *bak1-4* and *bak1-3serk4-1* (**D**) seedlings were treated with 100 nM of TAMRA-Pep1 for 10 s (pulse) and imaged after 40 min (chase). Homozygous seedlings for the triple mutants were selected based on phenotype. **(C, E)** Plasma membrane (PM) vs. intracellular fluorescence intensity measurements of the TAMRA-Pep1 uptake in **(B, D)**. Dots, red lines, and whiskers represent individual measurements, averages, and SD, respectively. Statistically significant differences were analyzed with one-way ANOVA followed by Tukey’s test. \*\*\*\**P*<0.0001; ns, not significant; *n*, number of cell analyzed. Epidermal cells of root meristematic zone were imaged (**A, B, D**). Scale bars, 10 µm (**A, B, D**).

**Supplemental Figure 3.**
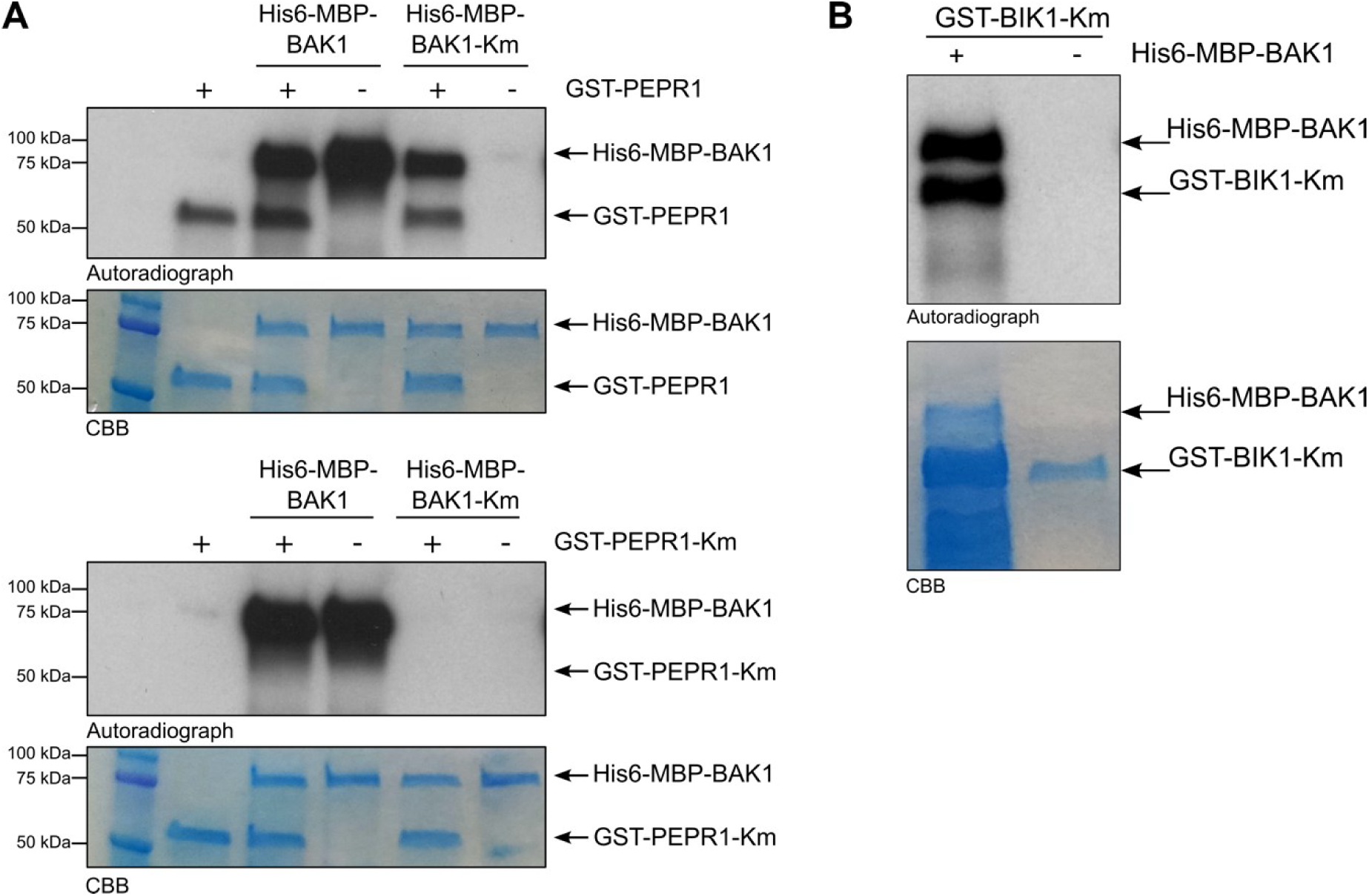
BAK1 cannot trans-phosphorylate PEPR1 *in vitro*. (**A**) The cytoplasmic domains of BAK1 and PEPR1, and their respective kinase-dead mutant (Km) versions of PEPR1-Km (K855E) and BAK1-Km (D416N) tagged with either glutathione S-transferase (GST) or His6-maltose-binding protein (MBP) were produced in bacteria. *In vitro* kinase assay performed with a single or combined recombinant proteins as indicated. CBB, Coomassie Brilliant Blue. (**B**) His6-MBP-BAK1 can phosphorylate the inactive BIK1 kinase (BIK1-Km). For the *in vitro* kinase assay a single or recombinant proteins together were used, as indicated. Protein loading was controlled by Coomassie Brilliant Blue (CBB).

**Supplemental Figure 4.**
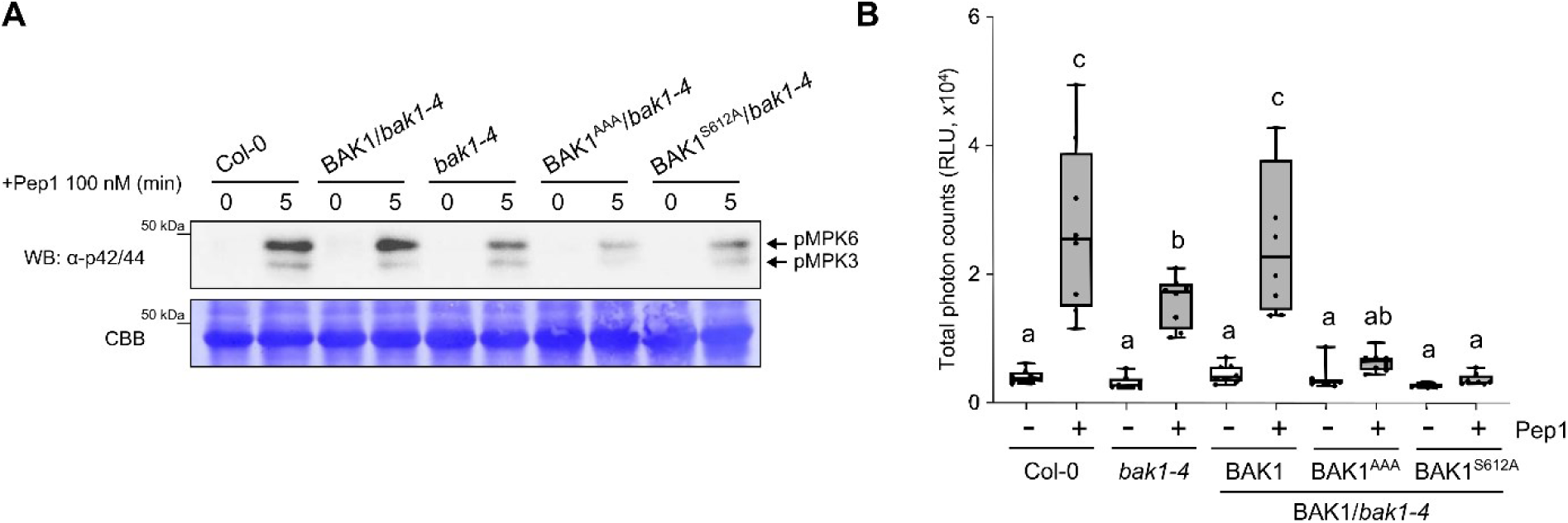
MAPK activation and ROS burst after Pep1 treatment were impaired in the BAK1^S612A^ and BAK1^AAA^ mutants. (**A**) Pep1-mediated MAPK activation was compromised in BAK1^S612A^ and BAK1^AAA^ mutants as shown in *bak1-4* mutant. 9-day-old seedlings of wild-type (Col-0), *pBAK1-BAK1*/*bak1-4*, *pBAK1-BAK1^S612A^/bak1-4*, and *pBAK1-BAK1*^AAA^/*bak1-4* were treated with 100 nM Pep1 and with mock (water) for 5 min. MAPK activation was analyzed with α-pERK1/2 antibodies (top), and protein loading is shown by CBB staining for RBC (bottom). (**B**) The BAK1^S612A^ and BAK1^AAA^ mutants showed compromised Pep1-induced ROS burst. Leaf discs (*n* = 8) from 4-week-old soil-grown plants were treated with or without 300 nM Pep1, and ROS production was measured as relative light units (RLUs) by a luminometer. Data represent mean ± SEM

**Supplemental Figure 5.**
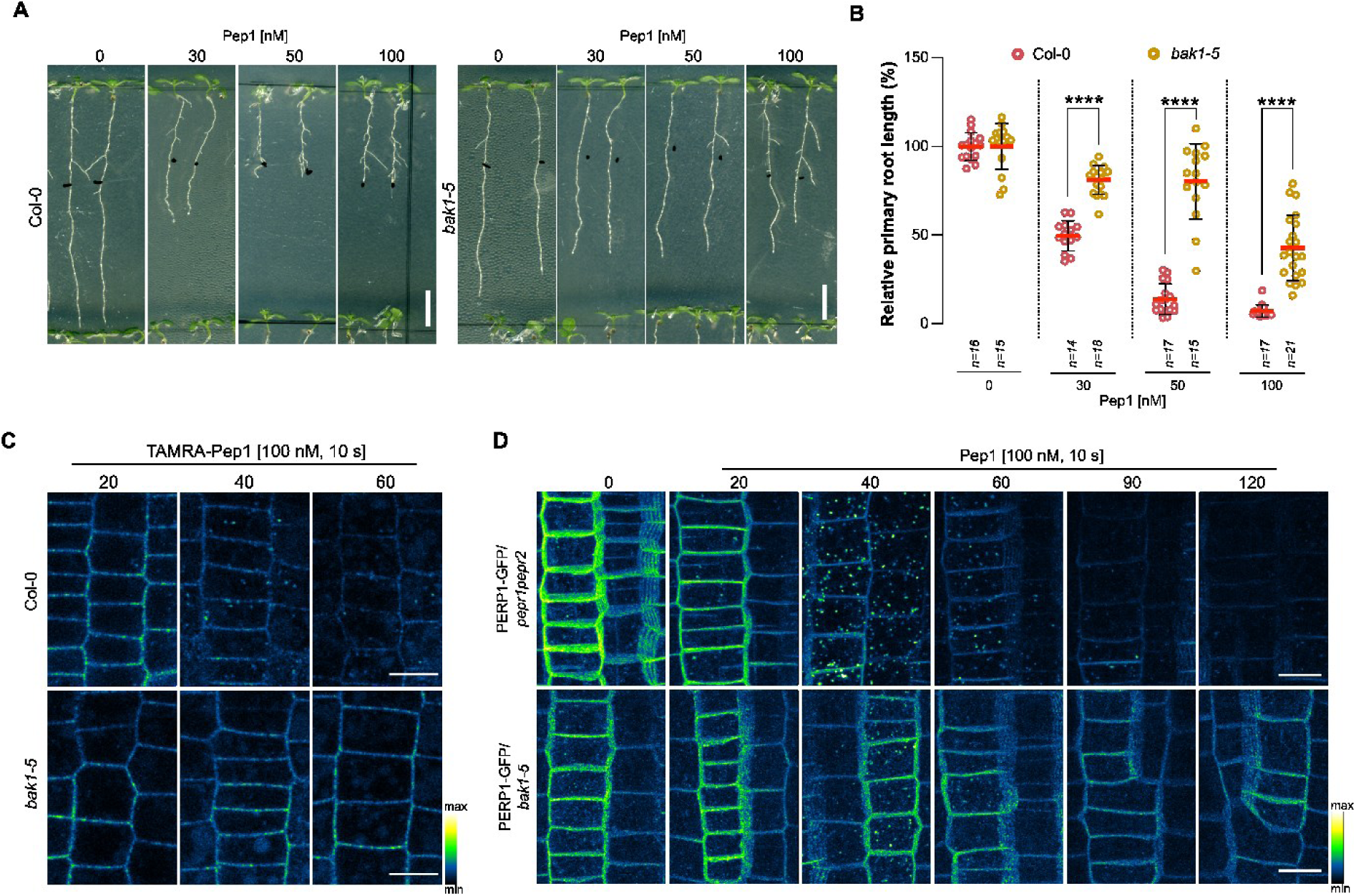
The hypoactive kinase BAK1-5 disrupts PEPR1 endocytosis and signaling. **(A)** Representative images of the root growth inhibition caused by different Pep1 concentrations. Seedlings of wild-type Col-0 and *bak1-5* were grown for 4 days on ½MS agar plates and transferred to ½MS agar media supplemented with different concentrations of Pep1 or water (mock) for 3 days. Scale bars, 10 mm. **(B)** Primary root length of seedlings in **(A)** presented as relative to the mock-treated control for each genotype. Dots, red lines, and whiskers represent individual measurements, averages, and SD, respectively. The two-way ANOVA followed by Tukey’s test was used to determine significant differences. \*\*\*\**P*<0.0001; *n*, number of seedlings analyzed. **(C, D)** Representative images of TAMRA-Pep1 root uptake **(C)** and PEPR1-GFP endocytosis **(D)** in the *bak1-5* mutant visualized by color-based fluorescence intensity coding. Five-day-old seedlings of wild-type Col-0 and *bak1-5* **(C)**, and *pRPS5A-PEPR1-GFP*/*pepr1pepr2* and *pRPS5A-PEPR1-GFP*/*bak1-5* were treated with 100 nM of TAMRA-Pep1 **(C)** and 100 nM Pep1 **(D)** for 10 s and imaged at indicated time points (chase). Epidermal cells of root meristematic zone were imaged. Images are maximum intensity Z-projections. Scale bars, 10 µm.

**Supplemental Figure 6.**
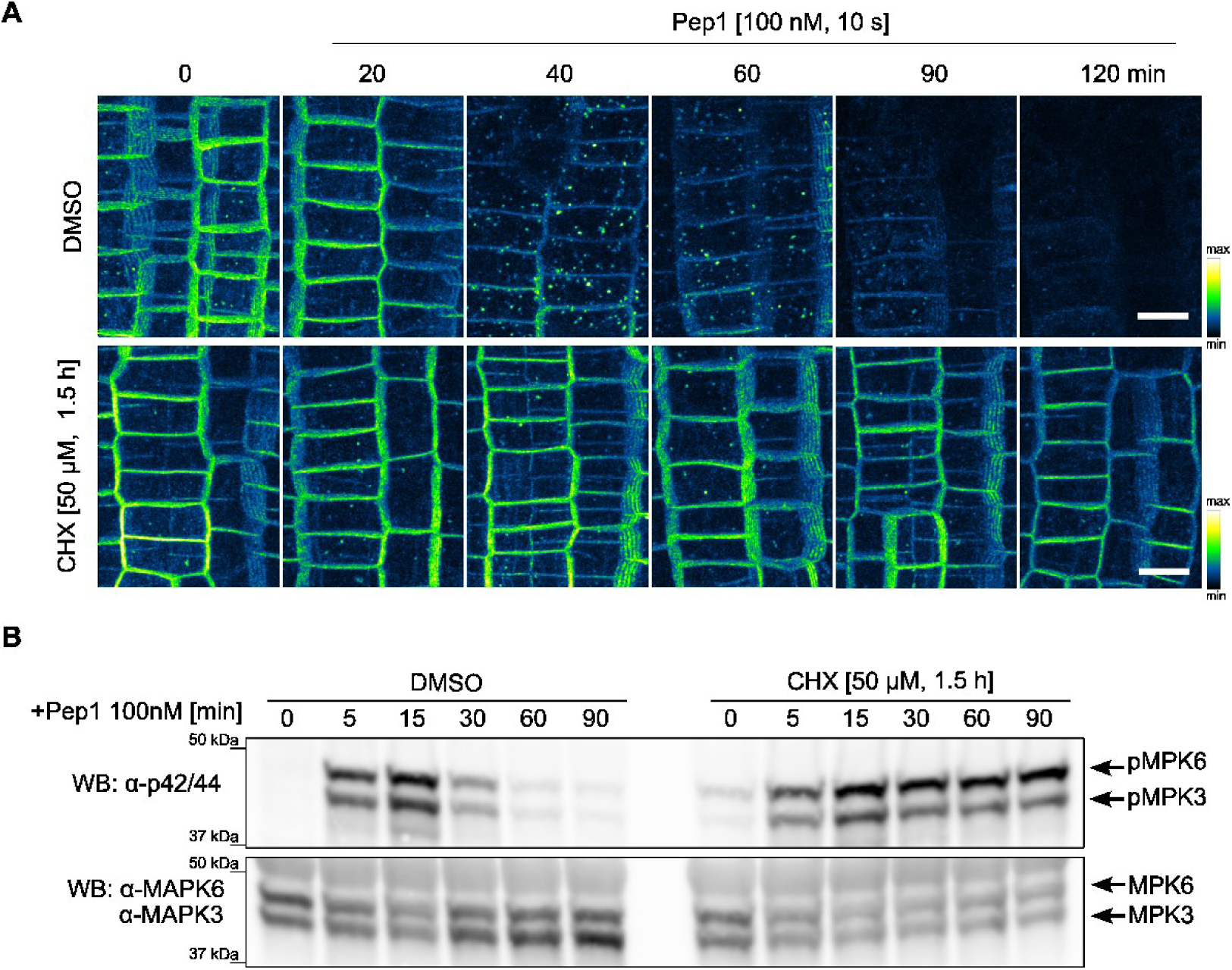
Cycloheximide (CHX) blocks PEPR1-GFP endocytosis and affects Pep1-triggered MAPK activation. **(A)** Representative images of PEPR1-GFP internalization visualized by color-based fluorescence intensity coding in the presence of CHX. Five-day-old *pRPS5A-PEPR1-GFP*/*pepr1pepr2* seedlings were pre-treated with DMSO (mock) and CHX (50 µM, 1.5 h) followed by application of 100 nM Pep1 for 10 s (pulse) and imaged at the indicated time points (chase). Epidermal cells of root meristematic zone were imaged. Images are maximum intensity Z-projections. Scale bars, 10 µm. **(B)** MAPK activation assay after Pep1 treatment in the presence of CHX. Five-day-old wild-type Col-0 seedlings were pre-treated with DMSO (mock) or CHX (50 µM, 1.5 h) followed by application of 100 nM Pep1 for the indicated time points. Phosphorylated MPK3 (pMPK3) and MPK6 (pMPK6) were detected by Western Blotting (WB) with α-p42/44 antibody (top). Protein loading of total MPK3 and MPK6 was detected with the α-MPK3 and α-MPK6 antibodies (bottom).

**Supplemental Figure 7.**
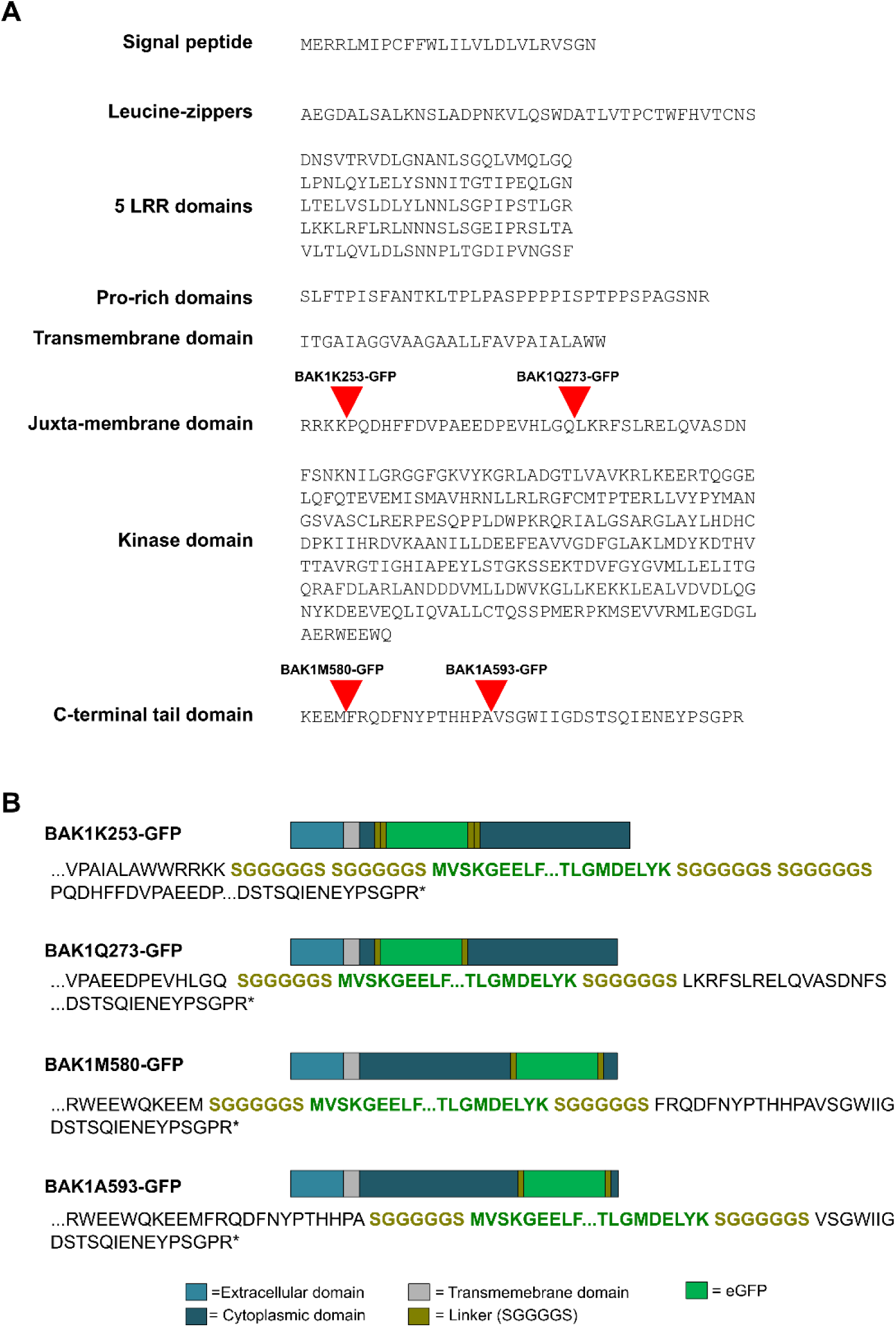
Schematic representation of the different constructs containing GFP-tagged BAK1. **(A)** Amino acid sequences of the different BAK1 protein domains. The positions of the GFP tags are indicated with red triangle. LRR, leucine-rich repeat. **(B)** Schematic representation of the tagged BAK1-GFP lines with the detailed amino acid sequences of the insertional junction regions.

**Supplemental Figure 8.**
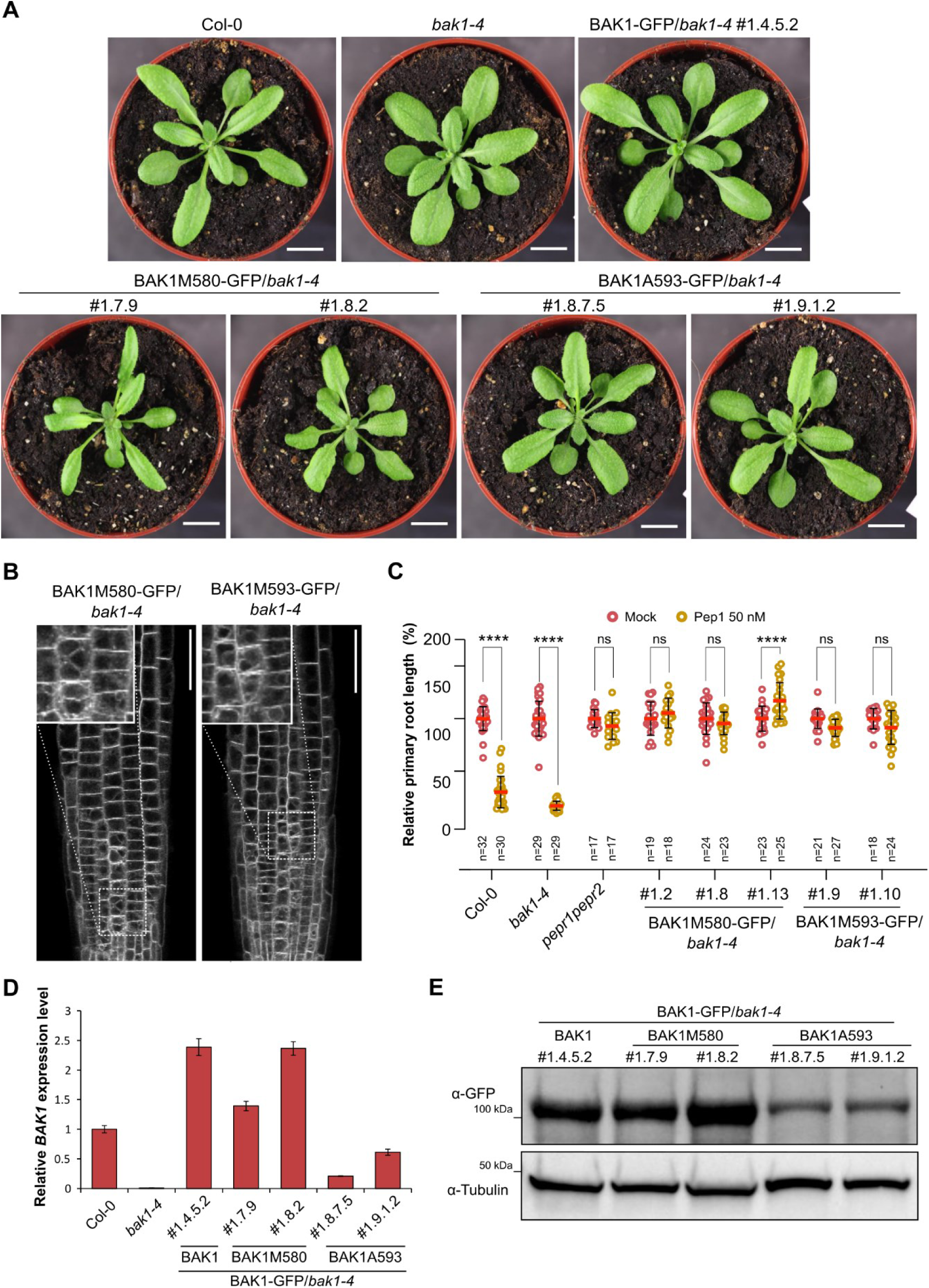
GFP insertional mutagenesis screen for functional BAK1-GFP-expressing Arabidopsis lines. **(A)** Representative images of 4-week-old T3 plants of the indicated genotypes. Scale bars, 1 cm. **(B)** Subcellular localization of BAK1-GFP in different GFP insertion lines. Root meristems of 5-day-old seedlings were imaged. Scale bars, 10 µm. **(C)** Relative primary root length of seedlings of the indicated genotypes grown for 4 days on ½MS agar plates and transferred to ½MS solid media supplemented with 50 nM Pep1 or water (mock) for 3 days. Dots, red lines, and whiskers represent individual measurements, averages, and SD, respectively. The two-way ANOVA followed by Tukey’s test was used to determine significant differences.\*\*\*\**P*<0.0001; ns, not significant; *n*, number of seedlings analyzed. **(D)** Real-time quantitative reverse transcription PCR (qRT-PCR) analysis of *BAK1*. Total RNA was isolated from 7-day-old seedlings of two independent transgenic lines. Transcript levels were normalized to *UBQ* gene with three technical replicates of each were quantified. Data are means ± SD **(E)** Total proteins isolated from 7-day-old seedlings and detected by Western Blotting (WB) with α-GFP antibodies to detect the BAK1-GFP. Numbers indicate individual transgenic lines. α-tubulin antibodies were used to equilibrate protein loading.

**Supplemental Figure 9.**
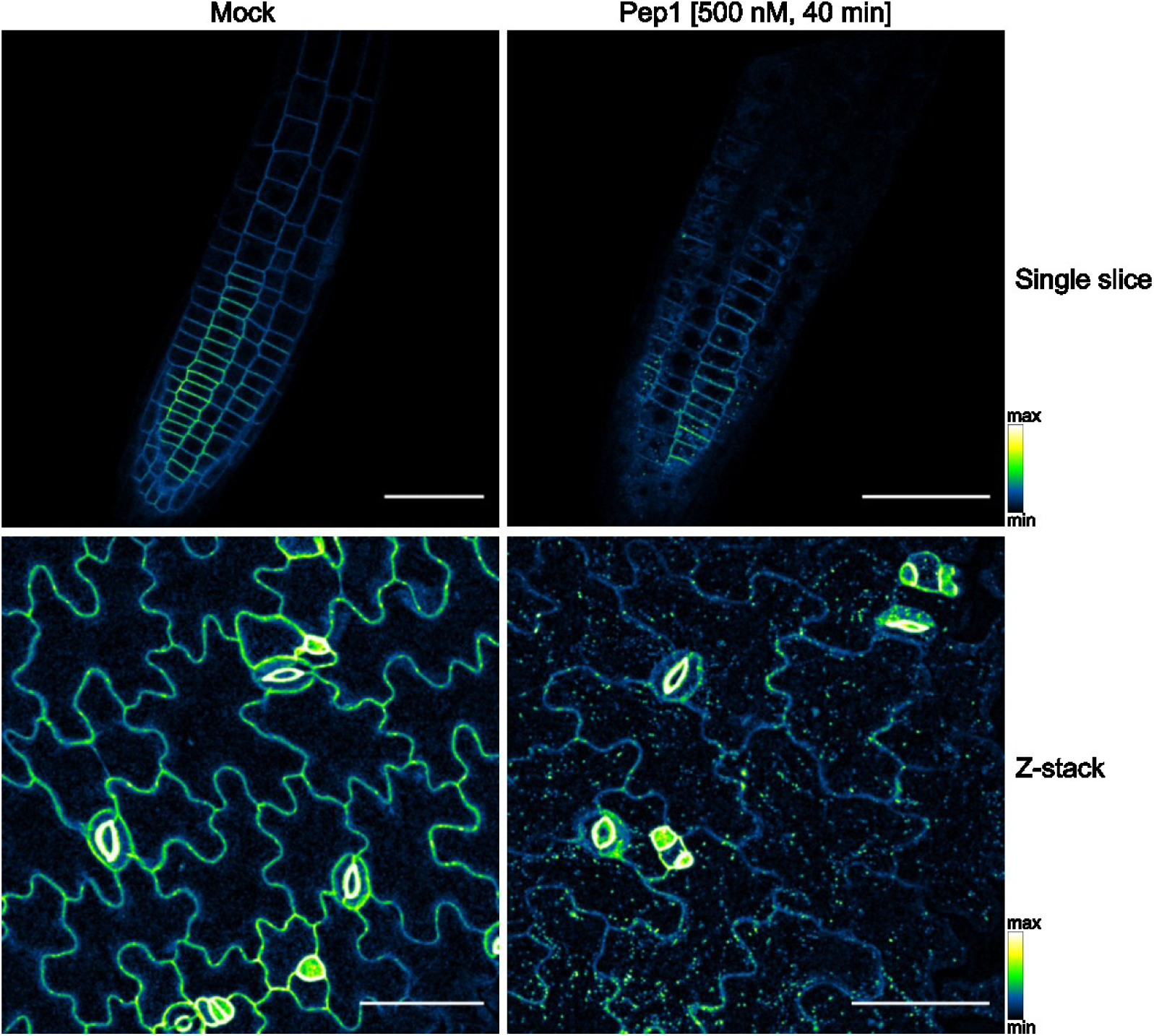
PEPR1-GFP endocytosis in Arabidopsis root and shoot. Representative images of PEPR1-GFP internalization visualized by color-based fluorescence intensity coding. Ten-day-old *pRPS5A-PEPR1-GFP*/*pepr1pepr2* seedlings were treated with water (mock) or 500 nM Pep1 for 40 min and root (top) and cotyledon (bottom) were imaged under confocal microscope. Images are maximum intensity Z-projections. Scale bars, 50 µm.

**Supplemental Table 1.**
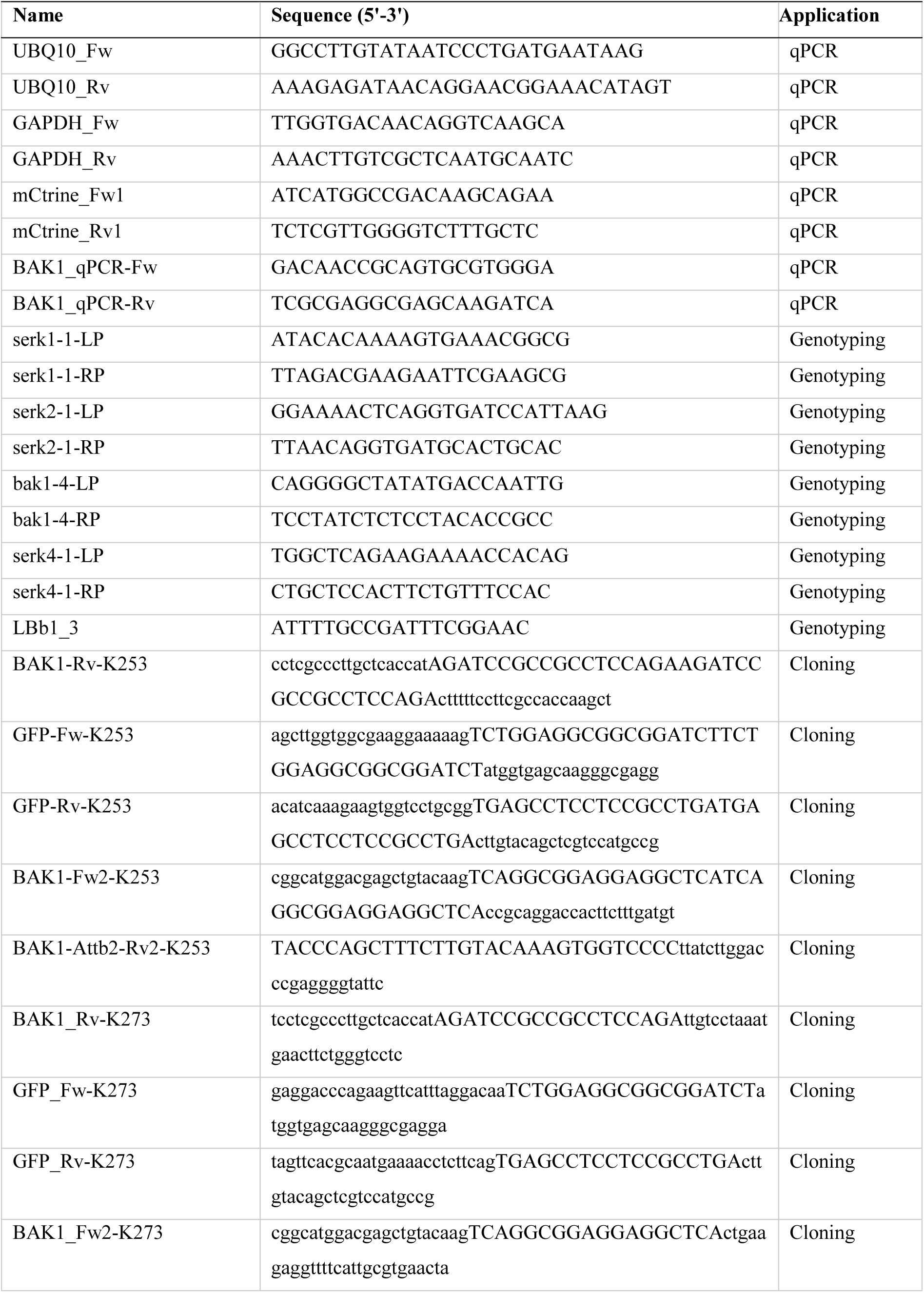
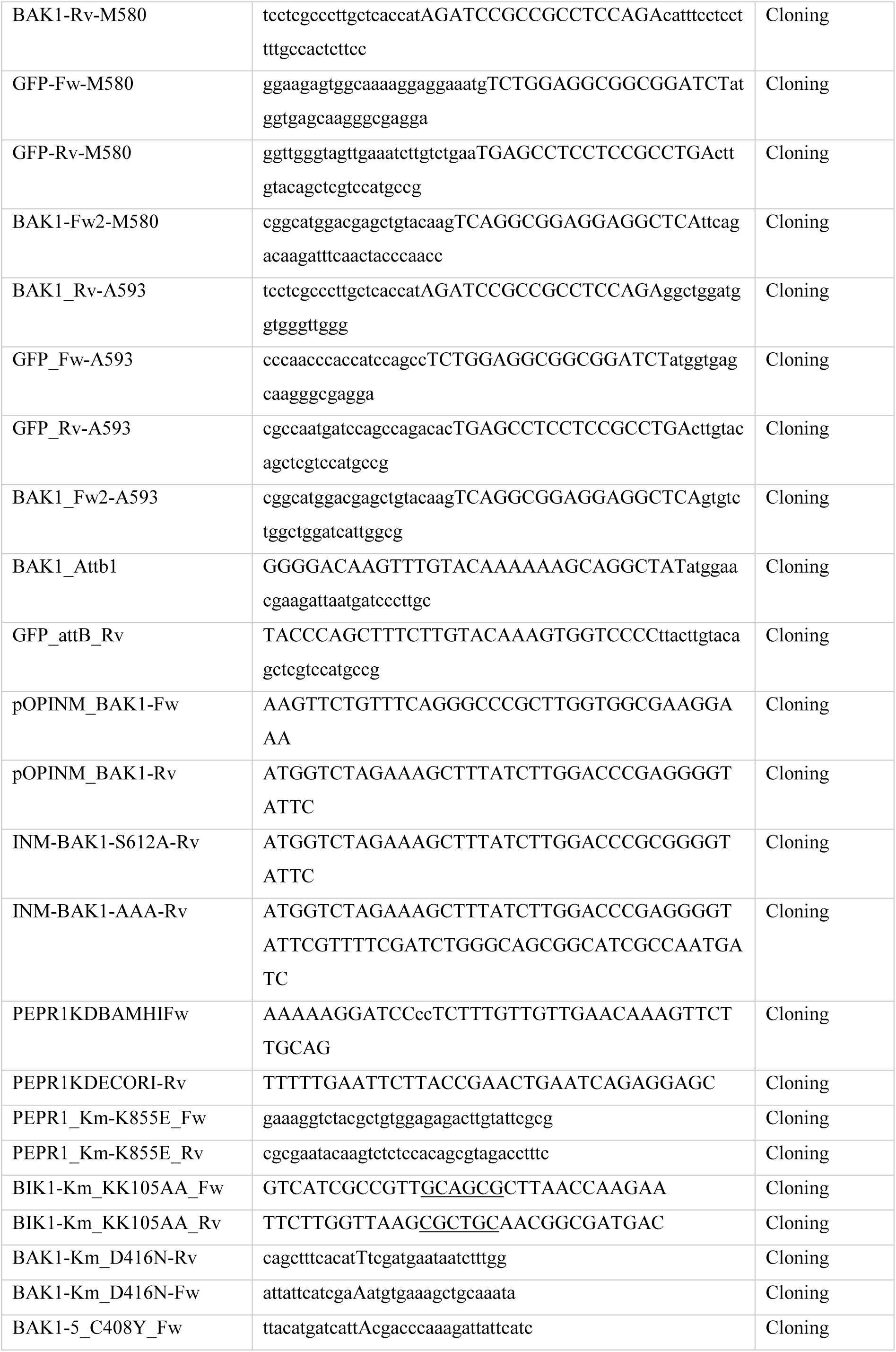
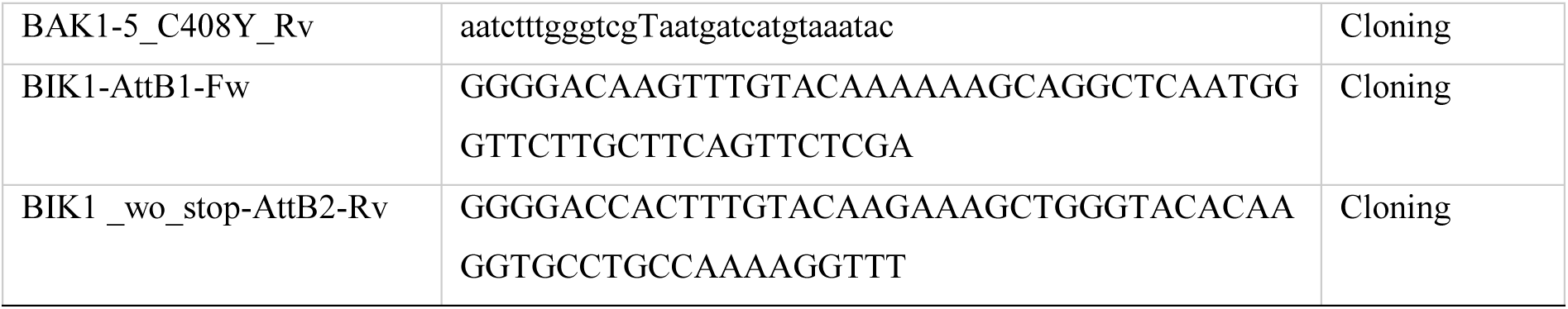
Primers used in this study.

## REFERENCES

Bartels S, Boller T (2015) Quo vadis, Pep? Plant elicitor peptides at the crossroads of immunity, stress, and development. J Exp Bot 66: 5183–5193

Bartels S, Lori M, Mbengue M, van Verk M, Klauser D, Hander T, Böni R, Robatzek S, Boller T (2013) The family of Peps and their precursors in *Arabidopsis*: differential expression and localization but similar induction of pattern-triggered immune responses. J Exp Bot 64: 5309–5321

Beck M, Zhou J, Faulkner C, MacLean D, Robatzek S (2012) Spatio-temporal cellular dynamics of the Arabidopsis flagellin receptor reveal activation status-dependent endosomal sorting Plant Cell 24: 4205–4219

Bender KW, Couto D, Kadota Y, Macho AP, Sklenar J, Derbyshire P, Bjornson M, DeFalco TA, Petriello A, Font Farre M et al. (2021) Activation loop phosphorylation of a non-RD receptor kinase initiates plant innate immune signaling. Proc Natl Acad Sci USA 118: e2108242118

Berrow NS, Alderton D, Sainsbury S, Nettleship J, Assenberg R, Rahman N, Stuart DI, Owens RJ (2007) A versatile ligation-independent cloning method suitable for high-throughput expression screening applications. Nucleic Acids Res 35: e45

Bücherl CA, van Esse GW, Kruis A, Luchtenberg J, Westphal AH, Aker J, van Hoek A, Albrecht C, Borst JW, de Vries SC (2013) Visualization of BRI1 and BAK1(SERK3) membrane receptor heterooligomers during brassinosteroid signaling Plant Physiol 162: 1911–1925

Chinchilla D, Zipfel C, Robatzek S, Kemmerling B, Nürnberger T, Jones JDG, Felix G, Boller T (2007) A flagellin-induced complex of the receptor FLS2 and BAK1 initiates plant defence. Nature 448: 497–500

Choi J, Tanaka K, Cao Y, Qi Y, Qiu J, Liang Y, Lee SY, Stacey G (2014) Identification of a plant receptor for extracellular ATP. Science 343: 290–294

Claus LAN, Savatin DV, Russinova E (2018) The crossroads of receptor-mediated signaling and endocytosis in plants. J Integr Plant Biol 60: 827–840

Clough SJ, Bent AF (1998) Floral dip: a simplified method for *Agrobacterium*-mediated transformation of *Arabidopsis thaliana*. Plant J 16: 735–743

Du J, Gao Y, Zhan Y, Zhang S, Wu Y, Xiao Y, Zou B, He K, Gou X, Li G et al. (2016) Nucleocytoplasmic trafficking is essential for BAK1- and BKK1-mediated cell-death control. Plant J 85: 520–531

Dubeaux G, Neveu J, Zelazny E, Vert G (2018) Metal sensing by the IRT1 transporter-receptor orchestrates its own degradation and plant metal nutrition. Mol Cell 69: 953–964

Dubeaux G, Vert G (2017) Zooming into plant ubiquitin-mediated endocytosis. Curr Opin Plant Biol 40: 56–62

Ekanayake G, Smith JM, Jones KB, Stiers HM, Robinson SJ, LaMontagne ED, Kostos PH, Cornish PV, Bednarek SY, Heese A (2021) DYNAMIN-RELATED PROTEIN DRP1A functions with DRP2B in plant growth, flg22-immune responses, and endocytosis. Plant Physiol 185: 1986–2002

Erwig J, Ghareeb H, Kopischke M, Hacke R, Matei A, Petutschnig E, Lipka V (2017) Chitin-induced and CHITIN ELICITOR RECEPTOR KINASE1 (CERK1) phosphorylation-dependent endocytosis of *Arabidopsis thaliana* LYSIN MOTIF-CONTAINING RECEPTOR-LIKE KINASE5 (LYK5). New Phytol 215: 382–396

Fan M, Wang M, Bai M-Y (2016) Diverse roles of SERK family genes in plant growth, development and defense response. Sci China Life Sci 59: 889–896

Geldner N, Anders N, Wolters H, Keicher J, Kornberger W, Muller P, Delbarre A, Ueda T, Nakano A, Jürgens G (2003) The *Arabidopsis* GNOM ARF-GEF mediates endosomal recycling, auxin transport, and auxin-dependent plant growth. Cell 112: 219–230

Hammond JW, Blasius TL, Soppina V, Cai D, Verhey KJ (2010) Autoinhibition of the kinesin-2 motor KIF17 via dual intramolecular mechanisms. J Cell Biol 189: 1013–1025

Hou S, Liu Z, Shen H, Wu D (2019) Damage-associated molecular pattern-triggered immunity in plants. Front Plant Sci 10: 646

Huffaker A, Dafoe NJ, Schmelz EA (2011) ZmPep1, an ortholog of Arabidopsis elicitor peptide 1, regulates maize innate immunity and enhances disease resistance. Plant Physiol 155: 1325–1338

Huffaker A, Pearce G, Veyrat N, Erb M, Turlings TCJ, Sartor R, Shen Z, Briggs SP, Vaughan MM, Alborn HT et al (2013) Plant elicitor peptides are conserved signals regulating direct and indirect antiherbivore defense. Proc Natl Acad Sci USA 110: 5707–5712

Hurst CH, Turnbull D, Myles SM, Leslie K, Keinath NF, Hemsley PA (2018) Variable effects of C-terminal fusions on FLS2 function: not all epitope tags are created equal. Plant Physiol 177: 522–531

Irani NG, Di Rubbo S, Mylle E, Van den Begin J, Schneider-Pizoń J, Hniliková J, Šiša M, Buyst D, Vilarrasa-Blasi J, Szatmári A-M et al (2012) Fluorescent castasterone reveals BRI1 signaling from the plasma membrane. Nat Chem Biol 8: 583–589

Irani NG, Di Rubbo S, Russinova E (2014) In vivo imaging of brassinosteroid endocytosis in Arabidopsis. Methods Mol Biol 1209: 107–117

Jaillais Y, Vert G (2016) Brassinosteroid signaling and BRI1 dynamics went underground. Curr Opin Plant Biol 33: 92–100

Kim SY, Xu Z-Y, Song K, Kim DH, Kang H, Reichardt I, Sohn EJ, Friml J, Juergens G, Hwang I (2013) Adaptor protein complex 2–mediated endocytosis is crucial for male reproductive organ development in *Arabidopsis*. Plant Cell 25: 2970–2985

Krol E, Mentzel T, Chinchilla D, Boller T, Felix G, Kemmerling B, Postel S, Arents M, Jeworutzki E, Al-Rasheid KAS et al (2010) Perception of the *Arabidopsis* danger signal peptide 1 involves the pattern recognition receptor *At*PEPR1 and its close homologue *At*PEPR2. J Biol Chem 285: 13471–13479

Li B, Ferreira MA, Huang M, Camargos LF, Yu X, Teixeira RM, Carpinetti PA, Mendes GC, Gouveia-Mageste BC, Liu C et al (2019) The receptor-like kinase NIK1 targets FLS2/BAK1 immune complex and inversely modulates antiviral and antibacterial immunity. Nat Commun 10: 4996

Lin W, Li B, Lu D, Chen S, Zhu N, He P, Shan L (2014) Tyrosine phosphorylation of protein kinase complex BAK1/BIK1 mediates Arabidopsis innate immunity. Proc Natl Acad Sci USA 111: 3632–3637

Liu D, Kumar R, Claus LAN, Johnson AJ, Siao W, Vanhoutte I, Wang P, Bender KW, Yperman K, Martins S et al (2020) Endocytosis of BRASSINOSTEROID INSENSITIVE1 is partly driven by a canonical Tyr-based motif. Plant Cell 32: 3598–3612

Liu J, Chen S, Chen L, Zhou Q, Wang M, Feng D, Li J-F, Wang J, Wang H-B, Liu B (2017) BIK1 cooperates with BAK1 to regulate constitutive immunity and cell death in *Arabidopsis*. J Integr Plant Biol 59: 234–239

Lin Y, Huang X, Li M, He P, Zhang Y (2016) Loss-of-function of *Arabidopsis* receptor-like kinase BIR1 activates cell death and defense responses mediated by BAK1 and SOBIR1. New Phytol 212: 637–645

Liu Z, Wu Y, Yang F, Zhang Y, Chen S, Xie Q, Tian X, Zhou J-M (2013) BIK1 interacts with PEPRs to mediate ethylene-induced immunity. Proc Natl Acad Sci USA 110: 6205–6210

Lozano-Durán R, Macho AP, Boutrot F, Segonzac C, Somssich IE, Zipfel C (2013) The transcriptional regulator BZR1 mediates trade-off between plant innate immunity and growth. eLife 2: e00983

Lu D, Lin W, Gao X, Wu S, Cheng C, Avila J, Heese A, Devarenne TP, He P, Shan L (2011) Direct ubiquitination of pattern recognition receptor FLS2 attenuates plant innate immunity. Science 332: 1439–1442

Lu D, Wu S, Gao X, Zhang Y, Shan L, He P (2010) A receptor-like cytoplasmic kinase, BIK1, associates with a flagellin receptor complex to initiate plant innate immunity. Proc Natl Acad Sci USA 107: 496–501

Ma X, Claus LAN, Leslie ME, Tao K, Wu Z, Liu J, Yu X, Li B, Zhou J, Savatin DV et al (2020) Ligand-induced monoubiquitination of BIK1 regulates plant immunity. Nature 581: 199–203

Martins S, Dohmann EMN, Cayrel A, Johnson A, Fischer W, Pojer F, Satiat-Jeunemaître B, Jaillais Y, Chory J, Geldner N et al (2015) Internalization and vacuolar targeting of the brassinosteroid hormone receptor BRI1 are regulated by ubiquitination. Nat Commun 6: 6151

Mbengue M, Bourdais G, Gervasi F, Beck M, Zhou J, Spallek T, Bartels S, Boller T, Ueda T, Kuhn H et al (2016) Clathrin-dependent endocytosis is required for immunity mediated by pattern recognition receptor kinases. Proc Natl Acad Sci USA 113: 11034–11039

Meng X, Chen X, Mang H, Liu C, Yu X, Gao X, Torii Keiko U, He P, Shan L (2015) Differential function of *Arabidopsis* SERK family receptor-like kinases in stomatal patterning. Curr Biol 25: 2361–2372

Mosesso N, Bläske T, Nagel M-K, Laumann M, Isono E (2019) Preparation of clathrin-coated vesicles from *Arabidopsis thaliana* seedlings. Front Plant Sci 9: 1972

Mühlenbeck H, Tsutsui Y, Lemmon MA, Bender KW, Zipfel C (2024) Allosteric activation of the co-receptor BAK1 by the EFR receptor kinase initiates immune signaling. eLife 12: RP92110

Ntoukakis V, Schwessinger B, Segonzac C, Zipfel C (2011) Cautionary notes on the use of C-terminal BAK1 fusion proteins for functional studies. Plant Cell 23: 3871–3878

Ortiz-Morea FA, Savatin DV, Dejonghe W, Kumar R, Luo Y, Adamowski M, Van den Begin J, Dressano K, Pereira de Oliveira G, Zhao X et al (2016) Danger-associated peptide signaling in *Arabidopsis* requires clathrin. Proc Natl Acad Sci USA 113: 11028–11033

Pavlos NJ, Friedman PA (2017) GPCR signaling and trafficking: The long and short of it. Trends Endocrinol Metab 28: 213–226

Pei D, Hua D, Deng J, Wang Z, Song C, Wang Y, Wang Y, Qi J, Kollist H, Yang S et al (2022) Phosphorylation of the plasma membrane H^+^-ATPase AHA2 by BAK1 is required for ABA-induced stomatal closure in Arabidopsis. Plant Cell 34: 2708–2729

Perraki A, DeFalco TA, Derbyshire P, Avila J, Séré D, Sklenar J, Qi X, Stransfeld L, Schwessinger B, Kadota Y et al (2018) Phosphocode-dependent functional dichotomy of a common co-receptor in plant signalling. Nature 561: 248–252

Robatzek S, Chinchilla D, Boller T (2006) Ligand-induced endocytosis of the pattern recognition receptor FLS2 in *Arabidopsis*. Genes Dev 20: 537–542

Russinova E, Borst J-W, Kwaaitaal M, Caño-Delgado A, Yin Y, Chory J, de Vries SC (2004) Heterodimerization and endocytosis of Arabidopsis brassinosteroid receptors BRI1 and AtSERK3 (BAK1). Plant Cell 16: 3216–3229

Schulze B, Mentzel T, Jehle AK, Mueller K, Beeler S, Boller T, Felix G, Chinchilla D (2010) Rapid heteromerization and phosphorylation of ligand-activated plant transmembrane receptors and their associated kinase BAK1. J Biol Chem 285: 9444–9451

Schwessinger B, Roux M, Kadota Y, Ntoukakis V, Sklenar J, Jones A, Zipfel C (2011) Phosphorylation-dependent differential regulation of plant growth, cell death, and innate immunity by the regulatory receptor-like kinase BAK1. PLoS Genet 7: e1002046

Sigismund S, Confalonieri S, Ciliberto A, Polo S, Scita G, Di Fiore PP (2012) Endocytosis and signaling: cell logistics shape the eukaryotic cell plan. Physiol Rev 92: 273–366

Song L, Shi Q-M, Yang X-H, Xu Z-H, Xue H-W (2009) Membrane steroid-binding protein 1 (MSBP1) negatively regulates brassinosteroid signaling by enhancing the endocytosis of BAK1. Cell Res 19: 864–876

Takano J, Tanaka M, Toyoda A, Miwa K, Kasai K, Fuji K, Onouchi H, Naito S, Fujiwara T (2010) Polar localization and degradation of *Arabidopsis* boron transporters through distinct trafficking pathways. Proc Natl Acad Sci USA 107: 5220–5225

Tang J, Han Z, Sun Y, Zhang H, Gong X, Chai J (2015) Structural basis for recognition of an endogenous peptide by the plant receptor kinase PEPR1. Cell Res 25: 110–120

Tunc-Ozdemir M, Li B, Jaiswal DK, Urano D, Jones AM, Torres MP (2017) Predicted functional implications of phosphorylation of regulator of G protein signaling protein in plants. Front Plant Sci 8: 1456

Vu LD, Xu X, Zhu T, Pan L, van Zanten M, de Jong D, Wang Y, Vanremoortele T, Locke AM, van de Cotte B et al. (2021) The membrane-localized protein kinase MAP4K4/TOT3 regulates thermomorphogenesis. Nat Commun 12: 2842.

Wang S, Yoshinari A, Shimada T, Hara-Nishimura I, Mitani-Ueno N, Feng Ma J, Naito S, Takano J (2017) Polar localization of the NIP5;1 boric acid channel is maintained by endocytosis and facilitates boron transport in Arabidopsis roots. Plant Cell 29: 824–842

Wang X, Kota U, He K, Blackburn K, Li J, Goshe MB, Huber SC, Clouse SD (2008) Sequential transphosphorylation of the BRI1/BAK1 receptor kinase complex impacts early events in brassinosteroid signaling. Dev Cell 15: 220–235

Wang Y, Li Z, Liu D, Xu J, Wei X, Yan L, Yang C, Lou Z, Shui W (2014) Assessment of BAK1 activity in different plant receptor-like kinase complexes by quantitative profiling of phosphorylation patterns. J Proteomics 108: 484–493

Wang Z-Y, Seto H, Fujioka S, Yoshida S, Chory J (2001) BRI1 is a critical component of a plasma-membrane receptor for plant steroids. Nature 410: 380–383

Willems P, Sterck L, Dard A, Huang J, De Smet I, Gevaert K, Van Breusegem F (2024) The Plant PTM Viewer 2.0: in-depth exploration of plant protein modification landscapes. J Exp Bot 75: 4611–4624

Wiśniewska J, Xu J, Seifertová D, Brewer PB, Růžička K, Blilou I, Rouquié D, Benková E, Scheres B, Friml J (2006) Polar PIN localization directs auxin flow in plants. Science 312: 883–883

Wu D, Liu Y, Xu F, Zhang Y (2018) Differential requirement of BAK1 C-terminal tail in development and immunity. J Integr Plant Biol 60: 270–275

Yamada K, Yamashita-Yamada M, Hirase T, Fujiwara T, Tsuda K, Hiruma K, Saijo Y (2016) Danger peptide receptor signaling in plants ensures basal immunity upon pathogen-induced depletion of BAK1. EMBO J 35: 46–61

Yamaguchi Y, Huffaker A (2011) Endogenous peptide elicitors in higher plants. Curr Opin Plant Biol 14: 351–357

Yamaguchi Y, Huffaker A, Bryan AC, Tax FE, Ryan CA (2010) PEPR2 is a second receptor for the Pep1 and Pep2 peptides and contributes to defense responses in *Arabidopsis*. Plant Cell 22: 508–522

Yin Y, Wang Z-Y, Mora-Garcia S, Li J, Yoshida S, Asami T, Chory J (2002) BES1 accumulates in the nucleus in response to brassinosteroids to regulate gene expression and promote stem elongation. Cell 109: 181–191

Yu X, Xie Y, Luo D, Liu H, de Oliveira MVV, Qi P, Kim S-I, Ortiz-Morea FA, Liu J, Chen Y et al (2023) A phospho-switch constrains BTL2-mediated phytocytokine signaling in plant immunity. Cell 186: 2329–2344

Zhang J, Li W, Xiang T, Liu Z, Laluk K, Ding X, Zou Y, Gao M, Zhang X, Chen S et al (2010) Receptor-like cytoplasmic kinases integrate signaling from multiple plant immune receptors and are targeted by a *Pseudomonas syringae* effector. Cell Host Microbe 7: 290–301

Zhou J, Liu D, Wang P, Ma X, Lin W, Chen S, Mishev K, Lu D, Kumar R, Vanhoutte I et al (2018) Regulation of *Arabidopsis* brassinosteroid receptor BRI1 endocytosis and degradation by plant U-box PUB12/PUB13-mediated ubiquitination. Proc Natl Acad Sci USA 115: E1906–E1915

